# 3D multimodal histological atlas and coordinate framework for the mouse brain and head

**DOI:** 10.1101/2025.04.30.650246

**Authors:** Daniel Tward, Patrick J. Flannery, Stephen Savoia, Christopher Mezias, Samik Banerjee, Max S. Richman, Brianna M. Lodato, Joseph R. O’Rourke, Somesh Balani, Ken Arima, Stuart D. Washington, Ricardo Coronado-Leija, Jiangyang Zhang, Partha P. Mitra

## Abstract

Brain reference atlases are essential for neuroscience experiments and data integration. However, histological atlases of the mouse brain, crucial in biomedical research, have not kept pace. Autofluorescence-based volumetric brain atlases are increasingly used but lack microscopic histological contrast, cytoarchitectonic information, corresponding MRI datasets, and often have truncated brainstems. Here, we present a multimodal, multiscale atlas of the laboratory mouse brain and head. The new reference brains include the whole head with consecutive Nissl and myelin serial section histology in three planes of section with 0.46 µm in-plane resolution, including intact brainstem, cranial nerves, and associated sensors and musculature. We provide reassembled histological volumes with 20mu isotropic resolution in stereotactic coordinates, determined using co-registered *in vivo* MRI and CT. In addition to conventional MRI contrasts, we provide diffusion MRI-based *in vivo* and *ex vivo* microstructural information, adding a valuable co-registered contrast modality that bridges MRI with cell-resolution histological data. We shift emphasis from compartmental annotations to stereotactic coordinates in the reference brains, offering a basis for evolving annotations over time and resolving conflicting neuroanatomical judgments by different experts. This new reference atlas facilitates integration of molecular cell type data and regional connectivity, serves as a model for similar atlases in other species, and sets a precedent for preserving extra-cranial nervous system structures.

## Introduction

Neuroscientific research relies heavily on reference atlases, which provide a common framework for studying brain structure and function. However, traditional reference atlases have fundamental technical and conceptual limitations. These include presenting the brain in isolation rather than in the context of the surrounding head and its associated nervous system structures including cranial nerves, muscles and sensory organs, truncating the olfactory bulb and brainstem, lacking multimodality in imaging methods, using idiosyncratic rather than standardized coordinate spaces, and disagreements among expert neuroanatomists on regional boundaries and naming hierarchies. The lack of standardized outputs can impede the neuroscientific enterprise at a basic level, as researchers may be unable to accurately compare and interpret data across studies ^1,2^.

To overcome these issues, we propose and implement a comprehensive approach. Instead of providing a traditional segmented reference atlas with paired nomenclature, we constructed a Reference Atlas Framework (RAF). This framework consists of a set of multimodal reference brains at multiple scales, all embedded within the same stereotaxic reference space. It includes versioned and editable segmentations overlaid on these brains and a set of computational tools that permit continuous refinement of annotations and nomenclatures. This paper presents the Brain Architecture Project (BAP) Mouse Head RAF, which includes a multimodal set of downloadable data volumes with multiple MRI and histological contrasts, presented in a stereotactic coordinate system derived from multiple *invivo* MRI and CT datasets. The brain is presented in the context of the whole head, allowing better preservation of the connecting points into the brain via the cranial nerves, as well as the sensors (sense organs) and actuators (muscles) of the head. Interactive online viewers and annotation tools enable the atlas annotations to be refined and versioned over time, while the coordinate system itself remains fixed.

Even the best available histological atlases for model organisms such as mice (e.g., the Allen Reference Atlas) have major limitations. One significant drawback is the absence of *in vivo* MRI or other advanced imaging methodologies for comprehensive atlas purposes ^3^. Furthermore, these atlases often exhibit truncation or incomplete representation of the brainstem. On the other hand, many existing MRI atlases (**Extended Data Table 3**) are outdated, featuring relatively limited MRI contrasts that can capture the fine anatomical details in the mouse brain for comparisons with histology. Consequently, there is a pressing need for newer and updated versions of these atlases to address these limitations and provide more accurate and comprehensive representations of the mouse brain.

To enhance current atlas methodologies, we have acquired a multimodal dataset, including CT scans, MRI (*in vivo*, *ex vivo*), and histology (Nissl, myelin) data from the same animal. We collected histological series using three sectioning planes (coronal, axial, sagittal) with 10um section thickness, preserving the skull to keep brainstem structures intact. This is an improvement over existing atlases, which typically use one sectioning plane (coronal) and 100um spacing. We have also included a rich set of MR contrasts, especially quantitative parameter maps from diffusion MRI data using advanced biophysical models^4^, designed to extract structural organization at the cellular level for comparison with histology. In addition to the full head datasets, we have collected three brain-only serial Nissl series in the three sectioning planes with better preservation of brainstem structures. In this paper, we cross-modally registered all datasets and used skull landmarks from *in vivo* MRI and CT to define the stereotactic coordinate system with origin at bregma. The resulting dataset is available for viewing through the Brain Architecture portal, and the multimodal data volumes are available for download.

In summary, there are currently two main categories of mouse brain atlases: those based on histology and those based on MRI. While histology-based atlases, such as the Paxinos and Franklin atlas ^5^, have been widely used as reference volumes for the mouse brain with cellular-level information, MRI-based atlases provide a 3D anatomical framework undisrupted by sectioning and guide *in vivo* examination. We derived a new reference volume that combines the advantages of both techniques. Our atlas provides high-resolution histology, *in vivo* and *ex vivo* MRI, and *ex vivo* CT datasets. By incorporating both histology and MRI data, our reference volume offers a more comprehensive view of the mouse brain and has the potential to advance research in neuroscience. By providing a more flexible and adaptable reference atlas and improving upon existing datasets, researchers will be better equipped to accurately compare and interpret data across studies. This could lead to new insights and discoveries in neuroscience, ultimately leading to better treatments and therapies for neurological and psychiatric disorders.

## Results and discussion

### A next generation, whole head adult mouse brain reference atlas

Current mouse brain atlas resources have both technical and conceptual limitations, which the presented reference space resource aims to address. State-of-the-art brain atlases typically focus on microscopy, such as the Paxinos & Franklin ^6^ and Allen Institute resources ^3,7,8^, or radiology, like the Johns Hopkins resource ^9^ (**Table 1**). Additionally, current microscopy or histology-based atlases frequently lack 3D reconstructed volumes ^6^ or are not mapped to a meaningful coordinate grid, such as a stereotaxic reference space ^8^ (**Table 1**). Furthermore, there is no existing microscopy or histology-based atlas that includes images in all 3 sectioning planes (**Table 1; Extended Data Table 2**). Another key limitation of all current histological atlases is the exclusion of structures related to the rest of the nervous system in the head, and histological atlases often significantly truncate the brainstem and the olfactory bulb. We provide a brief review of existing histological mouse brain atlases in **Supplemental section S2** (histological atlases) and tabulate these atlases, along with literature references, in **Extended Data Table 2**. Similarly, we tabulate existing MRI atlases in **Extended Data Table 3**.

**Table 1.**
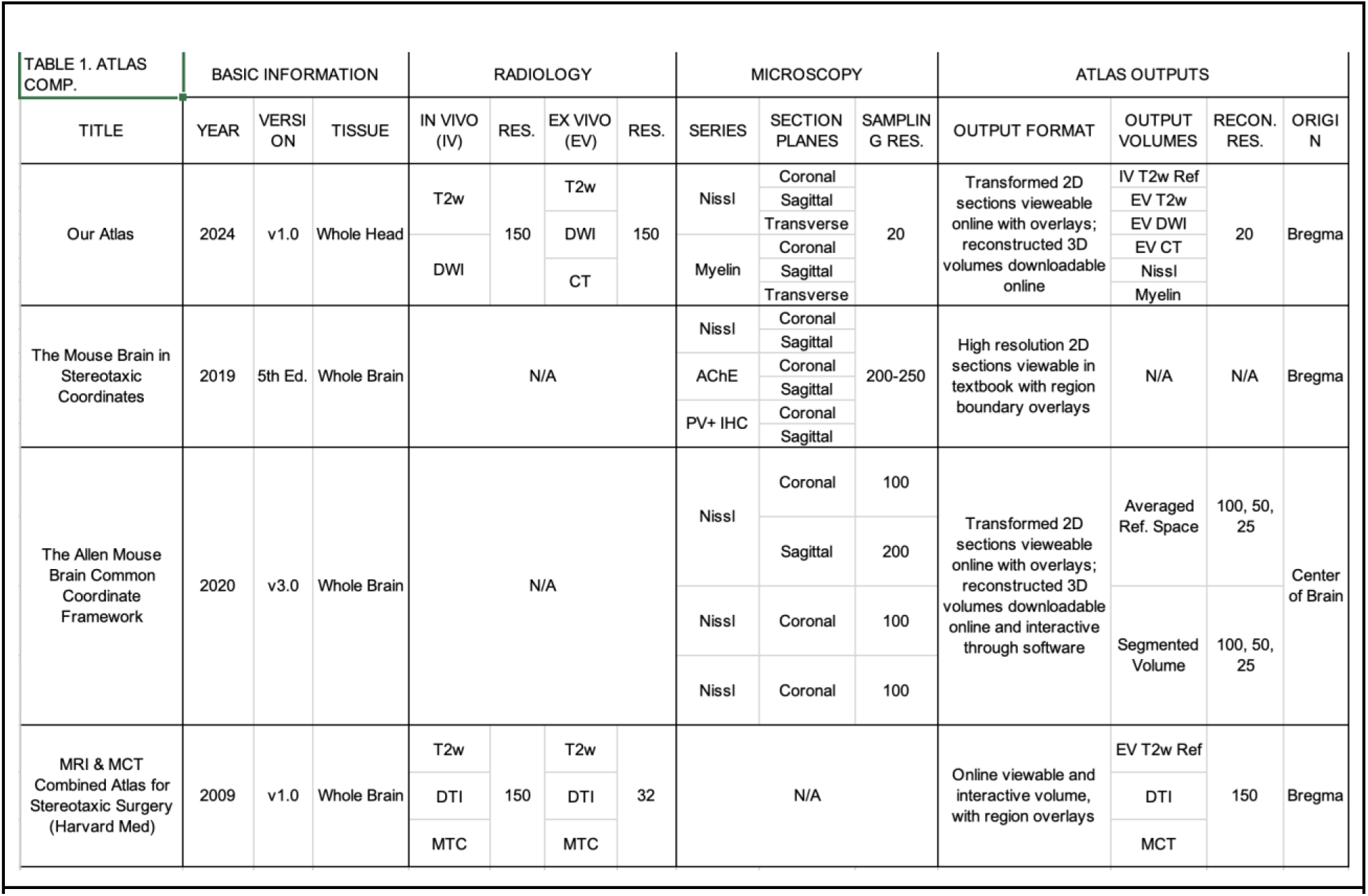
Comparison of the new and current field-standard mouse brain atlas resources. This table compares radiological and microscopy series sampling, resolution, atlas outputs and reference coordinate spaces. “N/A” indicates that a particular resource did not utilize a specific imaging modality.

**Table 2.** Atlas outputs are listed with columns noting whether and how they appear as both “Web Viewable” (https://www.brainarchitecture.org/bap-mouse-atlas/) and “Downloadable” outputs (https://data.brainarchitectureproject.org/pages/mouse). Usage of the atlas to register mouse brain data is demonstrated using data examples together with the associated code (https://data.brainarchitectureproject.org/pages/mouse)

| Table 2. Atlas Output | Web Viewable | Downloadable |
| --- | --- | --- |
| MRI series: (*in vivo*, *ex vivo*) x (T2W, Axial/Radial/Mean Diffusivity and Kurtosis, Fractional Anisotropy, | N/A | 20µm isotropic voxels<br>Reference Space volumes |
| Nissl Series: Head (Coronal, Sagittal, Transverse) | High resolution Section Images (0.46µm in plane resolution, 10µm thickness) | 20µm isotropic voxels<br>Reference Space volumes |
| Myelin Series: Head (Coronal, Sagittal, Transverse) | High resolution Section Images (0.46µm in plane resolution, 10µm thickness) | 20µm isotropic voxels<br>Reference Space volumes |
| Nissl Series: Brain (Coronal, Sagittal, Transverse) | High resolution Section Images (0.46µm in plane resolution, 20µm thickness) | 20µm isotropic voxels<br>Reference Space volumes |
| Brain Region Segmentations | Overlay on high resolution Nissl and myelin Images | Annotated volume |
| Curvilinear Atlas Coordinate Grid | Overlay on high resolution Nissl and myelin Images | N/A |
| Reference Atlas Coordinates | Retrievable on hover from high resolution Nissl and myelin images | N/A |

The present multimodal set of reference brains, along with auxiliary data objects (BAP Mouse Head RAF), offers several technical advantages. These can be best understood by summarizing our approach in constructing this atlas (**Fig. 1**). We begin by acquiring *in vivo* T2-weighted (T2w) and diffusion MRI, followed by *ex vivo* scans of the same subjects with an extensive array of MRI contrasts, as well as *ex vivo* CT (**Fig. 1a**). Next, we decalcify the skull and section the entire cranium to produce whole head Nissl and myelin series (**Fig. 1a**). From the outset, we employ both radiological and histological, as well as *in vivo* and *ex vivo* imaging modalities, making the RAF truly multimodal. Since all imaging, including Nissl and myelin histology, is performed on the whole head, we also capture intact out-of-brain features, such as reconstructable peripheral nerves (**Fig. 1d**). We then use the CT data to fit bregma and the tangent plane, allowing us to impose a stereotaxic origin and orientation onto our averaged *in vivo* T2w template volume, with which all other series will be registered (**Fig 1b**). To address variability in cranial shape across individuals, we take a Fréchet mean of coordinate systems established from individual heads. We provide histological and up-sampled radiological volumes, reconstructed at 20𝛍m isotropic resolution, with the stereotaxic coordinate grid using bregma as the origin (**Fig. 1c**). In addition to the three whole-head datasets, we also collected brain-only Nissl serial sections (at 20𝛍m section spacing and in-plane resolution of 0.46𝛍m) in the three cardinal planes, with better preservation of brainstem structures as well as the olfactory bulb.

**Fig. 1.**
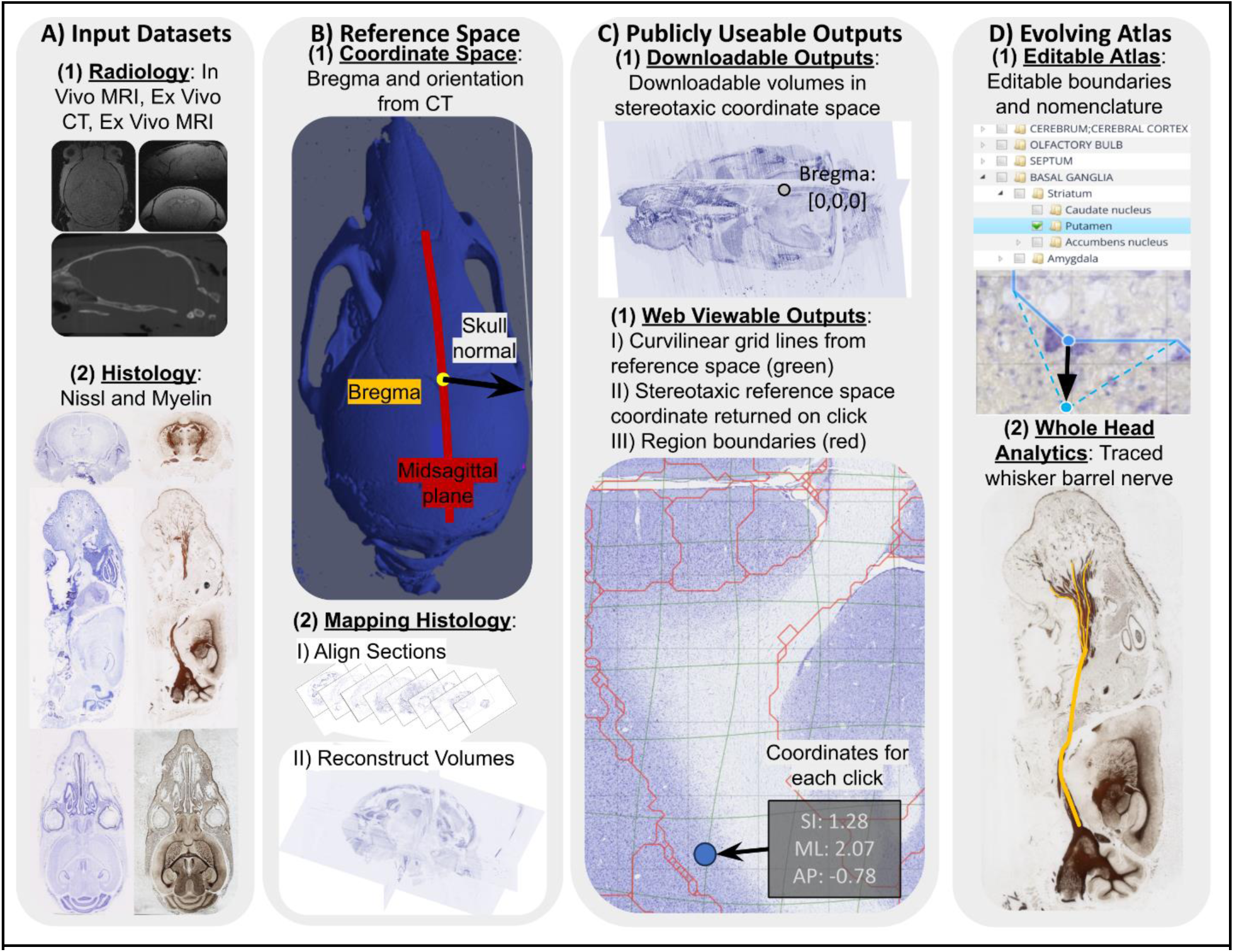
Summary of the pipeline for generating the next generation mouse common coordinate framework (CCFs) resource. **Column 1**: Whole-head *in vivo* and *ex vivo* MRI, CT and alternating Nissl and myelin histological series are acquired in the same animals. **Column 2**: A template volume is created by averaging *in vivo* MRI from 12 mice. The bregma location and tangent plane are determined by identifying the bregma location, skull normal vector, and symmetry plane from CT, which are then registered with the averaged template. Together, the averaged template from *in vivo* MRI and the averaged bregma location and tangent plane from CT form the basis of the atlas framework: reference volumes embedded within an *invivo* stereotactic coordinate system. Reconstructed volumes in reference space include *ex vivo* MRI series from three mice used for whole head histology, whole head Nissl and myelin histological series sectioned in coronal, transverse and sagittal planes, and an additional three brain-only serially sectioned Nissl datasets. Additionally, we provide corrected Allen Mouse Brain Atlas annotations registered with our reference space. **Column 3**: The atlas mappings of these datasets are available on the Brain Architecture web portal. This includes the averaged reference volume in stereotactic coordinate space, aligned 0.46 µm in-plane resolution histological sections with overlaid curvilinear coordinate grids and Allen region boundaries, and 20μm isotropic *ex vivo* MRI and histological volumes reconstructed in reference space. **Column 4**: We provide the starting point for an evolvable set of atlas annotations that we expect will be refined over time to include cranial nerves, sense organs and muscles in the mouse head, in addition to compartment annotations of the central brain.

We further disseminate the RAF interactively via the Brain Architecture portal and associated high-resolution 2D section viewer (https://www.brainarchitecture.org/bap-mouse-atlas/) and also provide isotropically resampled data volumes with multiple modalities and contrasts summarized in Table 2 (https://data.brainarchitectureproject.org/pages/mouse). In addition to displaying overlaid aligned adjacent Nissl and myelin sections, we have toggleable overlays for the curvilinear gridlines derived from the 3D stereotaxic reference space and region boundaries from the initial test segmentation ^8^, as well as the ability to query and return 3D atlas coordinates from 2D sections by mouse click (**Fig. 1c**). By creating a truly multimodal reference resource embedded within an *in vivo* stereotaxic coordinate space, we fill gaps in the technical capabilities of current atlases (**Table 1**).

Conceptually, the most widely used atlases, such as the Paxinos and Franklin ^6^ or Allen Institute ^3,7,8^ atlases, impose their particular version of region boundary segmentations onto their aligned histological and other microscopy sections, and in the case of the Allen Institute, their reconstructed volumes. These imposed image and volume segmentations derive from a pre-defined brain region ontological tree and label set. The BAP mouse head atlas eschews this top-down approach in favor of a flexible, evolvable and eventually user-defined set of compartment labels and region boundary segmentations. We have initiated a platform (**Fig. 1d**) that allows users to edit both compartment labels and region boundaries via a GUI built into the section viewer. We also link with a novel online image registration platform^10^ to enable users to register their radiological volumes or 2D microscopy with the averaged T2w *in vivo* reference space embedded within stereotaxic coordinates (**Fig. 1d**; **Fig 4**). We demonstrate the utility of this setup by registering example datasets with our reference space and provide the underlying data and code along with usage instructions (https://data.brainarchitectureproject.org/pages/mouse).

The BAP mouse head RAF represents an important conceptual shift away from fixed annotation-centric atlases towards evolvable reference atlases focused on reference brains and coordinate systems rather than specific compartment names and segmentations. In the subsequent sections, we describe both the underlying datasets and disseminated and interactive resources in greater detail.

### A new whole-head neurohistology reference space

An important feature of the BAP mouse head atlas is the whole head serial section histology with associated high-resolution digital microscopic images. Alternating series of neurohistological stains for cell bodies (Nissl stain) and myelinated axons (Gallyas myelin stain) were performed on serial sections in three cardinal planes in the three reference heads presented in the manuscript. The three reference brain-only volumes have serial Nissl stain in the three cardinal planes of section. High-resolution histological images from the three whole-head datasets are shown in coronal (**Fig. 2a**), transverse (**Fig. 2b**) and sagittal (**Fig. 2c**) sectioning planes. The three whole-head datasets were sectioned every 10𝛍m, with 20𝛍m spacing between sections of the same series (Nissl or myelin), and images were acquired with an in-plane resolution of 0.46𝛍m. The Nissl-only brain datasets were sectioned at 20𝛍m and imaged with an in-plane resolution of 0.46𝛍m.

**Fig. 2.**
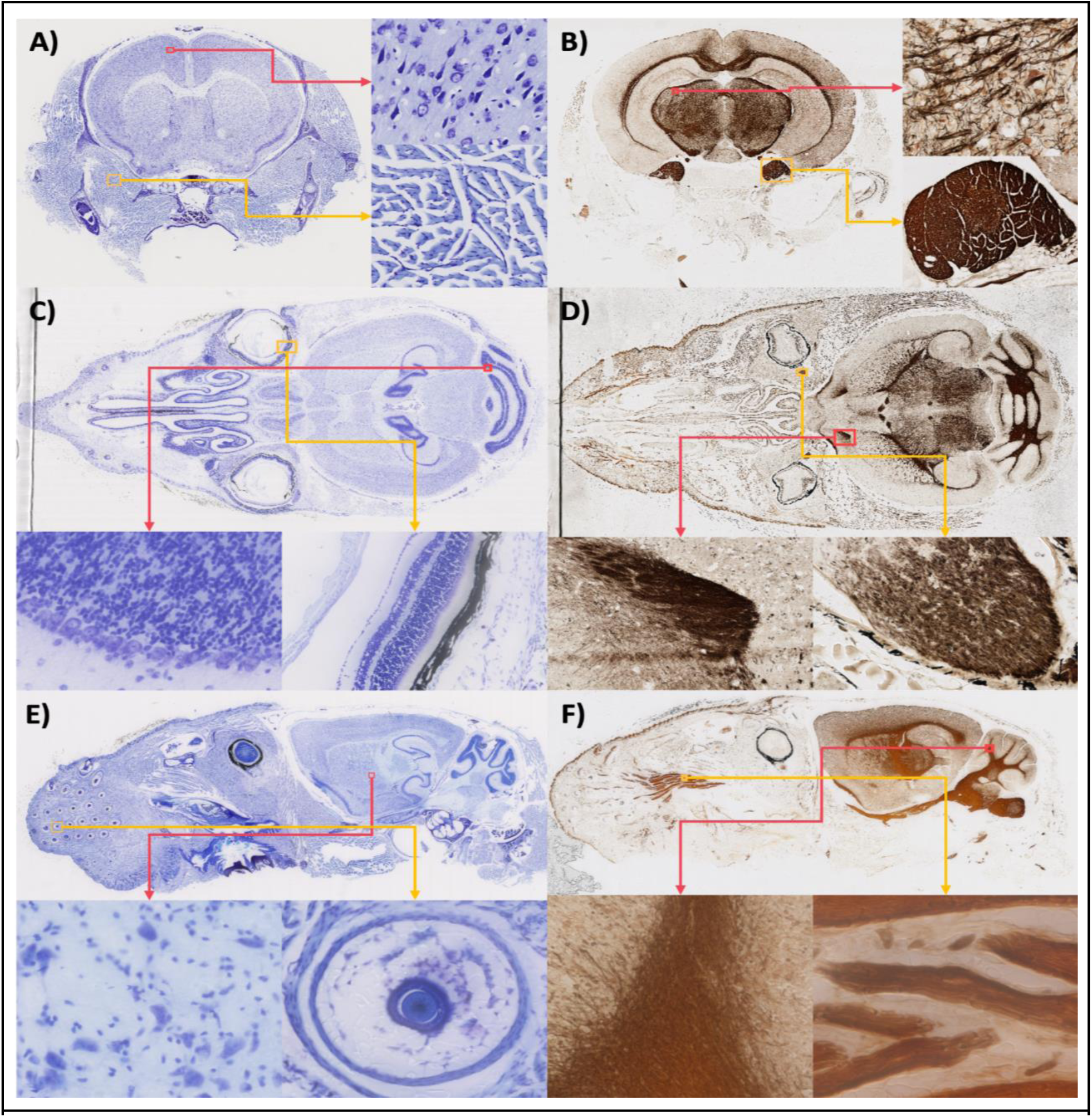
Example neurohistology (Nissl and myelin stain) data for the mouse head. We show example Nissl and myelin stain microscopy images of the mouse head in coronal (A-B), transverse (C-D) and sagittal (E-F) planes, captured at 0.46μm in-plane resolution. Each image includes zoomed-in cutouts demonstrating anatomical features both within and outside the brain. **A** Motor neurons (within brain) and muscle fibers (outside brain). **B** Fasciculated axons within the brain and a cross-section of a bundle of cranial nerves within the head below the brain. **C** Granule and Purkinje cell layers from the cerebellum within the brain, and optic features such as the retina and cornea outside the brain. **D** A frontal olfactory-related axon bundle within the brain, and parts of the optic nerve outside the brain. **E** Striatal medium spiny neurons and interneurons within the brain, and a cross-section of a whisker root outside the brain. **F** Cerebellar fiber bundles within the brain, and a section of the trigeminal nerve branching before innervation of the whisker field outside the brain.

We illustrate the histological stains in the head datasets by showing whole-section and cellular-level images (**Fig. 2a-c**), both within and outside the brain. We further demonstrate the advantages of sectioning the intact whole head in preserving peripheral nerves, including their entry/exit to/from the central brain and their extent within the head (**Fig 2d**). Additionally, cellular-level cytoarchitectonic definition is present across various structures in the head, such as muscles, retinal layers, and whisker barrels (**Fig. 2a-c**). Future segmentation of the whole-head histological datasets should permit quantitative delineation of such structures across the extent of the head with cellular resolution.

### Whole-head, combined *in vivo* and *ex vivo* multi-contrast MRI and CT

Most existing MRI-based atlases, with a few exceptions^9,11^, rely on high-resolution *ex vivo* mouse brain data to capture fine anatomical structures (e.g., ^12–14^, see **Supplementary Table 3** for a referenced list of existing MRI-based atlases together with key metadata). However, this approach limits the available tissue contrasts that are critical for understanding the diverse microstructural organization of the mouse brain. Our comprehensive MRI protocol addresses this limitation by including both *in vivo* and *ex vivo* T2-weighted scans for gross anatomical assessment, along with quantitative diffusion MRI parameter maps that indirectly measure tissue microstructural organization and integrity through empirical models of tissue water diffusion (**Fig. 3A**). We also incorporated advanced biophysical model parameters that reflect tissue microgeometry ^4^ (**Supplemental Section S3:** MRI Microgeometry Parameter Maps). The *ex vivo* dataset includes quantitative MR parameters sensitive to myelin and iron content^15^, such as longitudinal and transverse relaxation rates (R1 and R2*) and magnetization transfer saturation ratio (MTsat) (**Fig. 3B**). This dataset provides high-resolution 3D MRI data (0.1 mm isotropic resolution, interpolated to 0.05 mm isotropic) to facilitate precise alignment with histological sections.

**Fig. 3.**
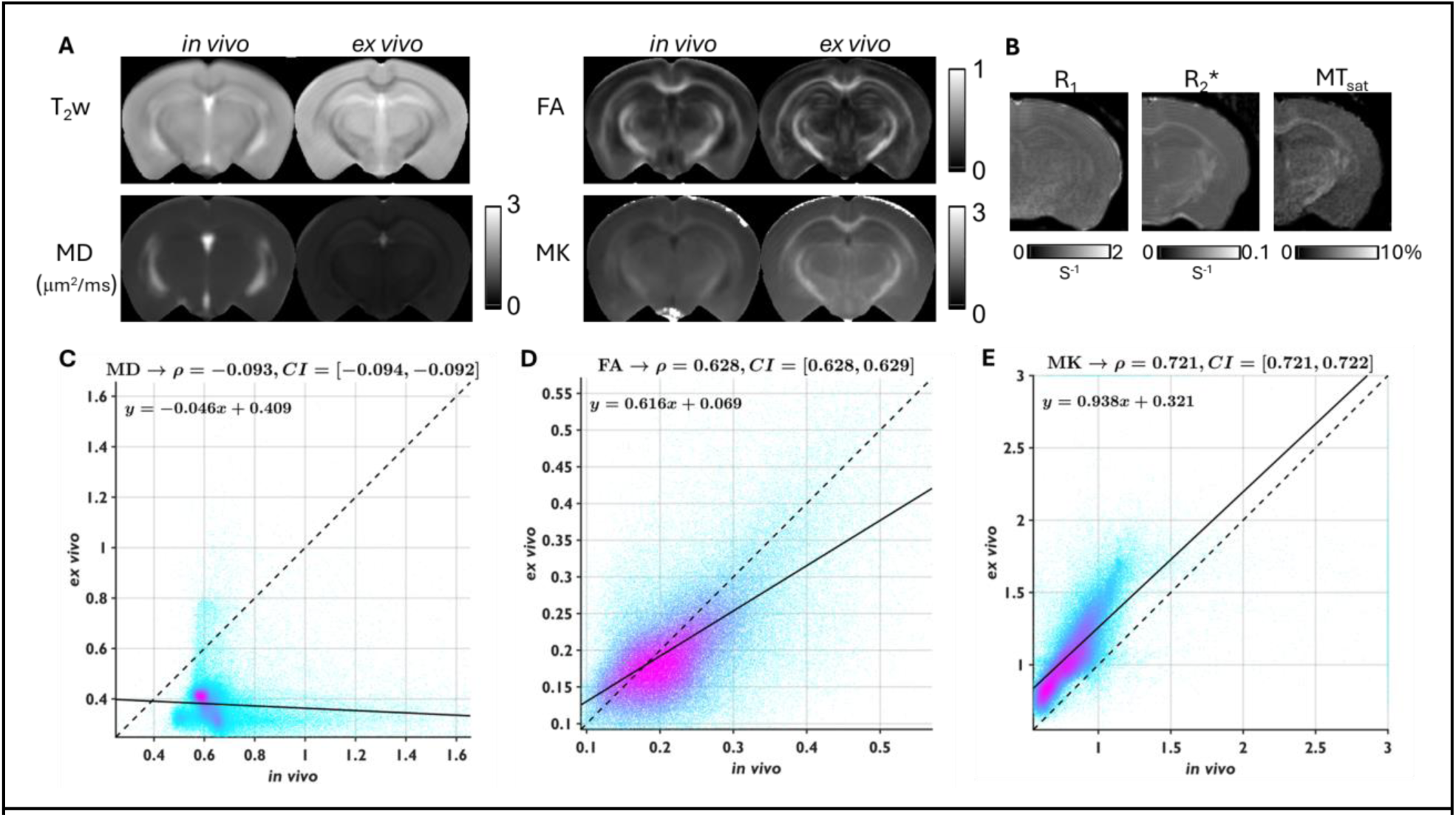
Representative *in vivo* and *ex vivo* modalities for the population-averaged female mouse brain. **A** Comparisons of *in vivo* and *ex vivo* T2-weighted (T2w) and diffusion MRI-derived maps, including fractional anisotropy (FA), mean diffusivity (MD) and mean kurtosis (MK). **B** Representative *ex vivo* maps of R1, R2* and MTsat, which are sensitive to myelinated white matter structures. **C-E** Voxel-wise comparisons between *in vivo* and *ex vivo* values, shown in scatter plots with Pearson correlation coefficient (𝜌) and 95% confidence interval (CI), characterizing differences in estimated MR parameters between the *in vivo* and *ex vivo* brain.

We are able to quantify the differences between the *in vivo* and *ex vivo* MRI datasets. Compared to *in vivo* MRI, which preserves the brain’s natural morphology critical for stereotaxic operations, *ex vivo* MRI data show postmortem alterations in brain morphology^9^ (e.g., reductions in ventricular size) and in estimated MR parameters, which reflect alterations in tissue microenvironments due to death, chemical fixation and temperature differences^16^. Our voxel-wise comparisons between *in vivo* and *ex vivo* MR parameters provide more comprehensive knowledge of these change than previous reports based on manually defined regions of interest^17^. These comparisons show reduced mean diffusivity (MD), slightly reduced fractional anisotropy (FA), increased mean kurtosis (MK) (**Fig. 3C-E)**, and related parameters **(Extended data Fig. 3)**. These changes likely reflect cell swelling as a result of the influx of water and sodium into the intracellular space after death. Parameters from the biophysical model (**Extended data Fig. 4**) revealed previously unreported subtle changes in tissue microstructure, including decreased axonal volume fraction (*f*) along with increased dispersion (p2), and an increase in the fraction of the ‘dot-compartment’ (*fiso*) ^18^, potentially corresponding to cell bodies.

### Mapping microscopy and radiology datasets to the BAP mouse head RAF

We provide a data and code example via GitHub to demonstrate how users can register their collected microscopy and radiology datasets with our new reference volumes (https://data.brainarchitectureproject.org/pages/mouse). This tool will allow users of the BAP mouse atlas to place their datasets within a stereotaxic coordinate space with bregma as the origin. It also enables users to embed brain-only datasets within our reference spaces, allowing data collected without the whole head to be placed within that context. The linked registration tool for BAP mouse atlas users preloads the same reconstructed downloadable reference volumes cited in Fig. 1 (histological and radiological).

To use the resource, users will need to input their collected microscopy or radiological datasets (**Fig. 4a**). They can also optionally change the default values of several parameters, such as the desired resolution for volumetric reconstruction and the optional co-registration of image stacks of serially histologically sections to create 3D input data. Once parameters are chosen, users can trigger the registration tool, Generative Diffeomorphic Mapping (GDM), whose algorithm is summarized in **Fig. 4b** and in recent work ^10^. Our approach to reference volume reconstruction is a generative probabilistic model, where a synthetic stack of 2D microscopy images is formed as a sequence of transforms of a 3D image, plus a noise model describing variability. Transforms include diffeomorphic spatial warping and contrast changes. Mapping to common coordinates is a maximum posterior estimation and enables the reconstruction of 3D and 2D datasets. We quantify tissue distortion with the derivative of spatial maps (morphometry) and account for scale changes when quantifying cell density. We support *in vivo* and *ex vivo* MRI, atlas annotations and multiple stained sections. This framework enabled us to jointly analyze multiple MRI contrasts and histology data, providing ground truth data for the evaluation of MR models.

Output file types will be consistent with those available with the BAP mouse head atlas datasets in **Fig. 1**. Each input dataset will be volumetrically reconstructed, registered and aligned as a stack of 2D sections at a user-defined resolution (**Fig. 4c**). Displacement fields, rotation matrices and Jacobian factors will also be saved per section, allowing users to retransform their data at different resolutions using the obtained registration outputs (**Fig. 4c**). Methodological information on how to use the registration outputs for transformation at arbitrary resolutions is detailed in the recent work ^10^ introducing the algorithm and package. Region boundary overlays from the user input set of ontological labels and segmented volume, and curvilinear coordinate gridlines from the BAP resource stereotaxic reference space, will also be returned as .json files (**Fig. 4c**).

By including a registration tool with the BAP atlas resource, we aim to facilitate community use of our unique multimodal resource. Allowing users to place their datasets in a stereotaxic coordinate grid, embedded within the context of the whole head, should enable new analyses and experiment planning, particularly for surgeries such as stereotaxic injections or electrode placement for electrophysiology.

**Fig. 4.**
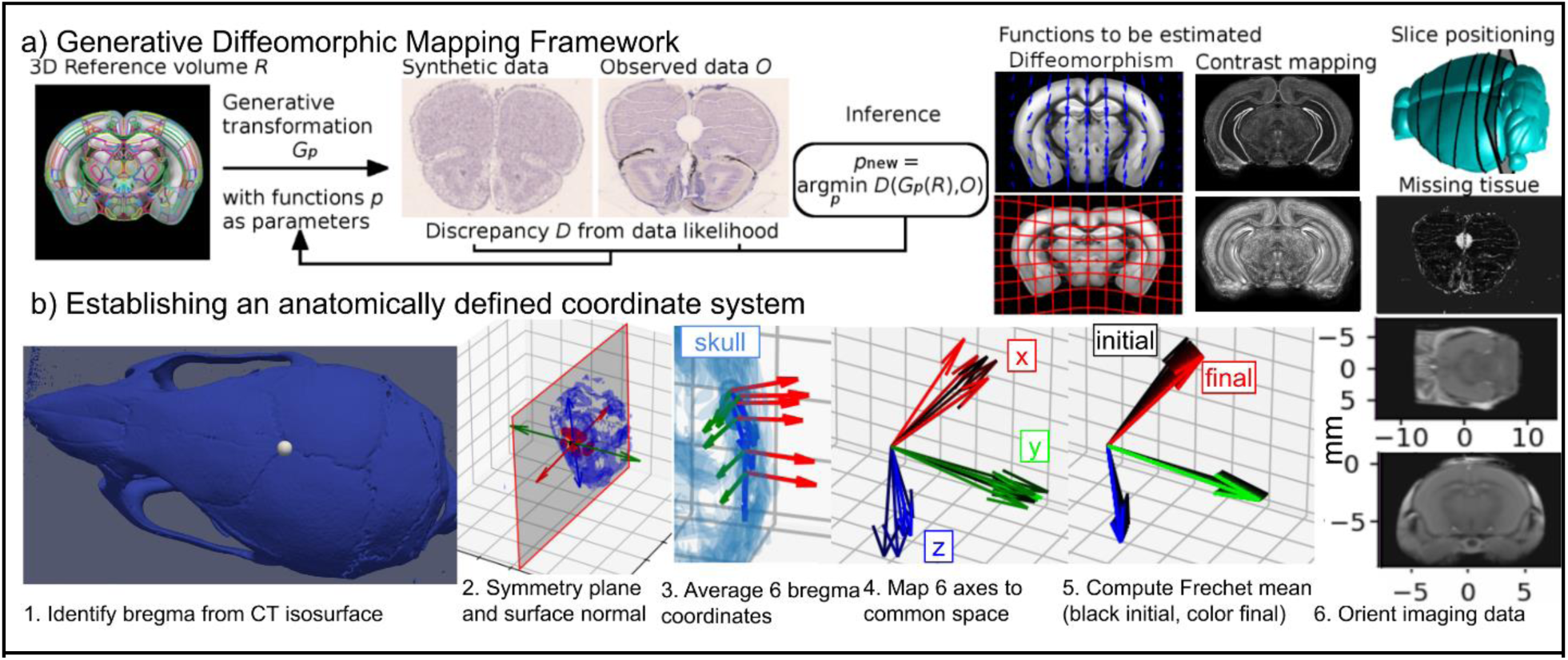
GDM registration algorithm for atlas mapping of radiological and histological data to the stereotactic mouse RAF. The GDM registration workflow is schematically shown in **(a**). Observed (input) datasets can include image series or volumes from the following modalities: *ex vivo* or *in vivo* MRI, histological brightfield or fluorescence microscopy, CT, and any volumetric data format. The reference brain volume, and optionally a segmented volume to obtain regional boundaries, can also be included. The algorithm estimates several unknown functions to synthesize the target dataset as a transformation of the reference volume. Outputs of the GDM registration pipeline include invertible transformations for computing high-resolution image transformations in any direction, high-resolution 2D microscopy and/or upsampled radiological image series, curvilinear gridlines representing the reference volume stereotactic coordinate system on 2D images, region boundary overlays for 2D images, and volumetric reconstructions of the input microscopy and/or radiological image series (outputs are shown in Fig. 1). (b) The procedure for identifying our coordinate system is described. b.1 On each CT dataset, the bregma point is identified on the skull. b.2-b.4. A symmetry plane and normal vector are estimated and mapped to a common space across six subjects, with an average bregma location estimated. b.5 An iterative Fréchet mean algorithm computes an average coordinate frame. The location of bregma was considerably variable along the AP axis of the skull (s.d.1.24mm), corresponding to individual biological variations of the Y-shaped bregma suture intersection. The Fréchet mean provides an average location and orientation suitable for our RAF. Note that our registration procedure for sample brains utilizes Nissl-based reference brain volumes on which this average coordinate system is superposed.

### An evolvable Reference Atlas Framework (RAF)

As noted in the first section of **Results** and **Fig. 1d**, a key conceptual advance of the BAP mouse atlas is its evolvability, allowing for user input and edits to nomenclature and segmentation. Instead of providing a traditional reference atlas, we constructed an evolvable Reference Atlas Framework, consisting of (i) a set of multimodal reference brains, (ii) versioned and editable segmentations overlayed on these brains, and (iii) online tools that enable a community-based effort to continuously refine the annotations and nomenclatures (**Fig. 1d**).

### Mapping single cell spatial transcriptomic data in BAP reference coordinates

To enhance the utility of our RAF for the neuroscience community, we have included spatial coordinates for each cell in the recently published Allen Brain Cell Atlas spatial single cell transcriptome ^19^. Using our registration algorithm, we computed a transformation between our reference space and the Allen Mouse CCF and applied this transformation to cell data accessed from the ABC atlas’ Python API. Cell IDs with coordinates in our reference space are available for download (https://data.brainarchitectureproject.org/pages/mouse) and can be used in conjunction with other cell-specific information provided in the ABC atlas. The spatial distribution of six cell types is shown in our RAF in **Fig. 5** and the spatial distribution of 35 cell types are shown in **Extended Data** Figure 6. The ability to superpose the spatial transcriptomics data with the three sectioning planes of light microscopic histological data in the BAP mouse atlas will help connect classical histological atlases to the new information available from the ongoing molecularly based cell-atlas efforts for the mouse.

**Figure 5.**
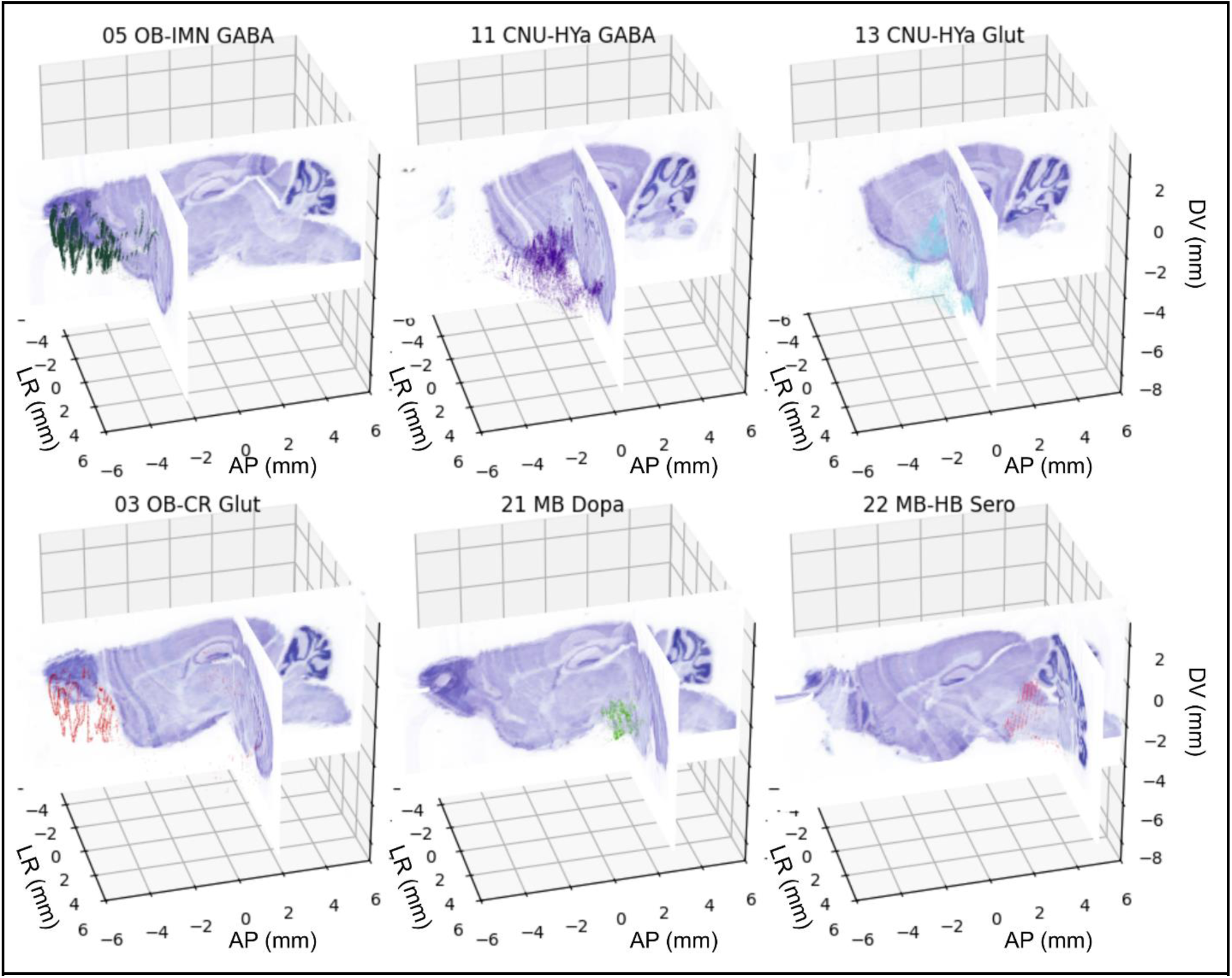
Spatial distributions of 6 selected cell types (see plot titles) from the ABC dataset, with coordinates mapped into our coordinate system. A 3D reconstructed Nissl reference brain dataset is shown in our coordinate system. Coronal and sagittal sections are chosen such that that 90% of cells appear in front of the slices. The visualization illustrates the intact brainstem present in the Nissl reference volume. Spatial distributions of a larger set of cell types are shown in **Extended Data** Figure 6.

### Nissl reference series for the mouse brain with better brainstem preservation

Existing histological reference atlases, including the one from the Allen Institute, do not have good preservation of the brainstem structures, resulting in less detailed annotations. To address this, our RAF includes three Nissl-only consecutive series of 20um thickness in the three cardinal planes of section (**Fig. 6**) with intact brainstem as well as olfactory bulb. The preservation of brainstem structures, with continuity into the cervical sections of the spinal cord, is demonstrated in Fig. 5, showing good preservation of cytoarchitectural features. We provide the corresponding reconstructed volumes together with our RAF resource, and offer the corresponding high-resolution series for online viewing and annotation (https://www.brainarchitecture.org/bap-mouse-atlas/). We believe that this will be a valuable dataset for studies involving non-coronally sectioned mouse brains.

**Fig. 6.**
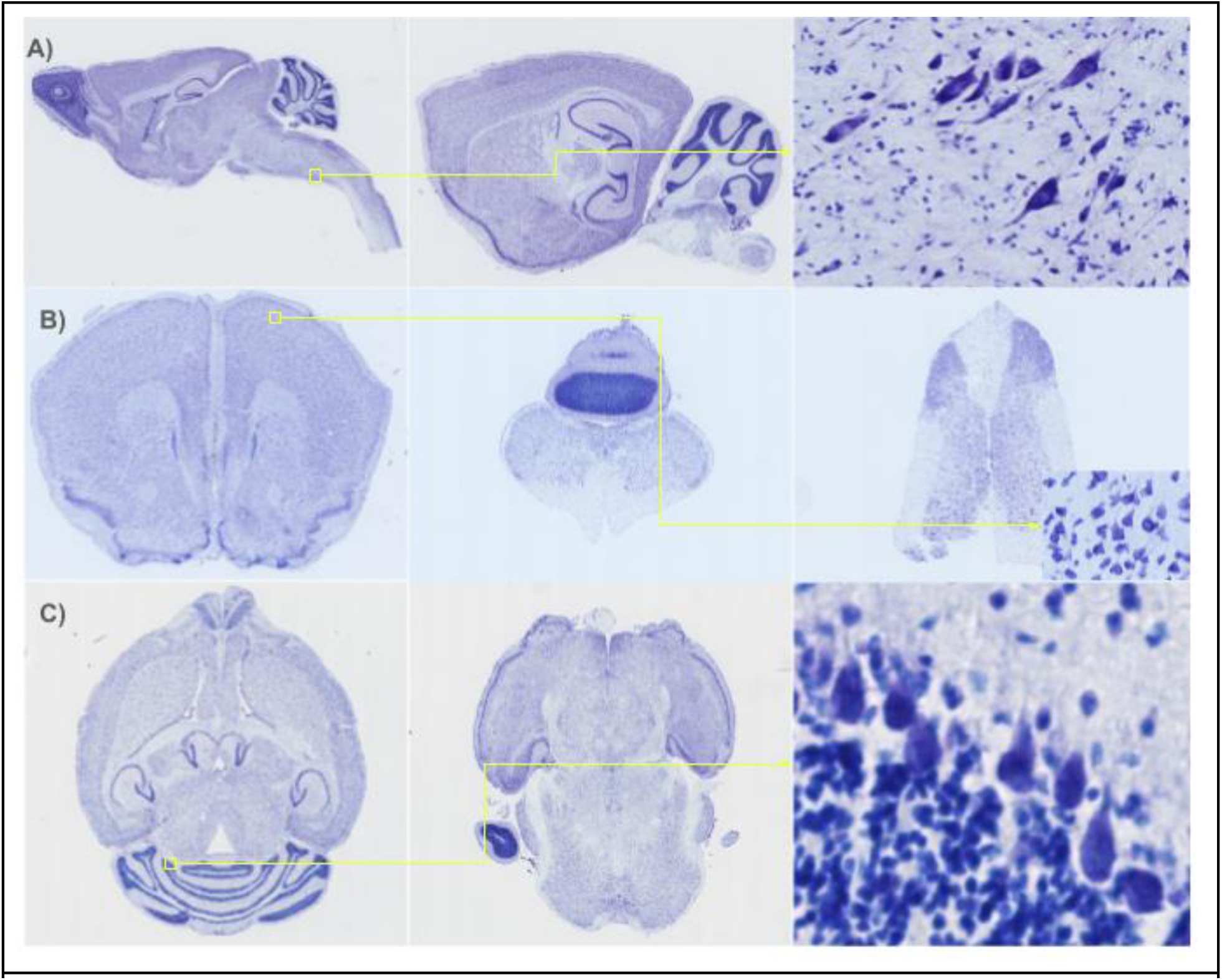
Nissl-only reference series for the mouse brain with no skull included but preserving brainstem structures with continuity into cervical spinal cord segments are provided in (a) sagittal, (b) coronal and (c) transverse planes. Serial 20um thick sections were Nissl (Thionin) stained and imaged at 0.46µm in-plane resolution. Zoomed-in cutouts show cellular resolution, allowing for the distinction of morphological types. Shown are (a) examples of medial and lateral sagittal sections and deep brainstem neurons, (b) example forebrain, brainstem and spinal cord coronal sections, as well as of neurons in layer 3 of the primary motor area, and (c) examples of medial and inferior transverse sections, as well as Purkinje and granular cells in the cerebellum.

### Segmented volume ID Reassignment and Integer Precision Remapping

The brain region segmented volumes used in the BAP mouse head RAF are derived from version 3 of the AIBS mouse atlas. Although there are only approximately 700 unique region IDs in the AIBS mouse atlas, the integer ID numerical values span a wide range, necessitating a high bit precision. Additionally, the original atlas did not separate homologous regions in the left and right hemispheres. To generate unique region IDs for the right hemisphere and avoid conflicts with existing IDs, higher values were used, requiring even greater bit precision. This approach results in large file sizes for annotated volumes and compatibility issues with standard volume viewing software. To address these issues, we regenerated all region IDs for the left hemisphere to range from 1 to 700 and adjusted the IDs for homologous regions in the right hemisphere by adding a fixed value. This allowed us to significantly reduce file sizes and ensure compatibility with standard volume viewing packages by using a lower bit precision format (see **Supplementary Section 5** for further details).

### Segmented Volume “Bubble Correction”

The brain region segmented volumes used in the BAP mouse head RAF are derived from version 3 of the AIBS mouse atlas. Due to registration and interpolation artifacts, these volumes contain bubbles, or isolated voxels with region IDs that do not match their neighbors. In the BAP mouse head RAF, we corrected these bubbles to eliminate these artifacts. First, we defined continuity of region IDs as voxels with the same ID assignment having touching faces. We set a threshold of five or fewer continuous voxels with the same ID assignment to constitute a bubble. We then identified all bubbles for each region ID and reassigned each voxel in each bubble to the same ID as the majority vote of its neighboring voxels, following a principle similar to nearest neighbor interpolation. This process was repeated until all voxels within bubbles were reassigned. Any remaining bubbles missed by the heuristic algorithm were manually corrected using Blender3D. This algorithm corrected approximately 99.5% of all voxels within bubbles and reduced the number of bubbles from almost 28,000 to just over 150, a reduction of over two orders of magnitude.

A Matlab package for generating these outputs, and the bubble-corrected, ID remapped reference volumes are provided in (https://data.brainarchitectureproject.org/pages/mouse). Further details are provided in **Supplementary Section 5 and Extended Data Fig. 5**.

## Conclusions

In this paper we have attempted to advance the state of the art in mouse neuroscience by introducing the BAP mouse head Reference Atlas Framework, consisting of a multimodal, multiscale co-registered set of data spanning in vivo and ex vivo MRI with multiple contrasts, ex vivo CT, as well as dense serial section histology of the whole mouse head with high resolution images of alternating Nissl and myelin-stained sections in Sagittal, Coronal and Transverse planes. In addition, we provide a set of brain-only serial section Nissl-stained data sets also in the three cardinal planes with better preservation of brainstem and olfactory bulb structures than is present in the literature.

We switch emphasis from a fixed set of atlas annotations to high information content underlying reference brains, together with an editable layer of annotations supplemented by online tools. To provide a starting point of the annotations, we provide an initial annotation volume which rectifies multiple issues with the current compartmental annotations available from the Allen mouse brain atlas including a very large number of isolated voxels. We provide data and code to enable users to register their own brain data sets to our new RAF.

To connect the RAF to ongoing spatial transcriptomics-based mouse brain atlases, we mapped the spatial coordinates of transcriptomically-typed single cells from the ABC atlas into our reference atlas framework. This will help bridge the new spatial transcriptomics-based atlases with the more established histological atlases which show cytoarchitectonic contrast that neuroanatomists have considered gold standards for atlas mapping of experimental data sets.

We carefully established a stereotactic coordinate system using same-animal CT scans, based on skull landmarks that are used in experimental neuroscience to place electrodes or injections in vivo. We present serial-section histology of the whole mouse head, to the best of our knowledge never presented previously, and which we hope will set a standard of preserving nervous system related structures outside of the central brain and permit better contextualization and understanding of central brain circuitry which has so far been analyzed largely in isolation from these closely related structures in the head.

## Material & Methods

### Subject Demographics & Colony Facilities

Mouse brains were collected at two institutions: Cold Spring Harbor Laboratory and New York University (NYU). Sectioning was performed in three planes: coronal, sagittal and transverse. All samples underwent histological processing, with serial section Nissl stain or alternating section Nissl and myelin stain. The head samples underwent skull decalcification. The list of brains used in this study and associated metadata are shown in **Extended Data Table 1**.

All C5BL/6J mice were sacrificed at P56 and radiologically imaged, if applicable. All experimental procedures were approved by the Animal Use and Care Committees at Cold Spring Harbor Laboratory and the New York University Grossman School of Medicine. See **Supplementary Section S4: Methods** for details.

### MRI/CT Image Series Acquisition Process

The same animals were scanned using computed tomography (CT) and magnetic resonance imaging (MRI), both *in vivo* and *ex vivo,* as listed in Supplementary Table 1.

*In vivo* MRI: Animals were anesthetized using isoflurane and imaged on a horizontal 7T MR scanner (Bruker Biospin, Billerica, MA, USA) using a 72-mm conventional circularly polarized birdcage radiofrequency resonator for homogeneous transmission and a four-channel receive-only phased array CryoProbe (CRP) for high sensitivity. Multi-slice T2-weighted images were acquired with an echo time (TE)/repetition time (TR) of 30/3000 ms, 4 signal averages, echo train length (ETL) of 8, an in-plane resolution of 0.125x0.125 mm^2^, and 33 axial slices with 0.5 mm thickness. Co-registered diffusion-weighted MR imaging (dMRI) was performed using a 4-segment echo planar imaging sequence with a diffusion gradient duration (𝛿)/diffusion time (𝛥) of 5/15 ms, 30 directions, and diffusion-weightings of 1000, 3000 and 5000 s/mm^2^. The total imaging time was less than 30 minutes for each mouse.

*Ex vivo* MRI: After perfusion fixation with 4% PFA, mouse brains were prepared as described in^20^. Multi-slice T2-weighted MRI of the entire mouse head was acquired first. For the brain, multi-slice T2-weighted and dMRI data were acquired using the same protocols as the *in vivo* MRI, with diffusion-weightings extended to 9000 s/mm^2^ due to approximately 50% reduction in tissue water diffusivity in *ex vivo* specimens compared to live mouse brains. High-resolution 3D *ex vivo* diffusion MRI datasets were acquired using a modified 3D diffusion-weighted gradient-and-spin-echo (DW-GRASE) sequence^21^ with a spatial resolution of 0.1x0.1x0.1 mm^3^, 𝛿/𝛥=5/15 ms, 60 directions, and diffusion-weightings of 5000 s/mm^2^. Additionally, quantitative magnetization transfer data were acquired using a 3D gradient echo sequence with the following parameters: TE/TR=2.1/45 ms, 12 echos with echo time spacing of 2.34 ms, 4 signal averages, offset frequency of 5 kHz, and a spatial resolution of 0.1x0.1x0.1 mm^3^.

dMRI images were pre-processed using the Designer^22^ toolbox (https://github.com/NYU-DiffusionMRI/DESIGNER-v2), including denoising^23^, Gibbs-ringing removal^24^, registration^25^, and Rician bias correction^26^. Using the *cumulant expansion*^27^ representation, we estimated^28^ the diffusion (D) and kurtosis (K) tensors that account for the Gaussian and non-Gaussian components of the diffusion process in the tissue. From D and K, several parameter maps were computed, such as principal diffusion direction, fractional anisotropy, axial, radial and mean diffusivities, and axial, radial and mean kurtosis.

*Ex vivo* CT: CT images were acquired using X-CUBE (Molecubes Inc.). The X-ray source was a tungsten anode with a focal spot size of 50 µm, filtered by a 0.8 mm aluminum filter. CT images were acquired using the following parameters: 360-degree rotation, 720 exposures, binning: 1x1; axial FOV: 37.4 mm, 4 averages, time per projection/gap time: 167 ms. The resolution of the reconstructed images was 0.05 mm isotropic.

Additional CT and MRI data, including 2D and 3D FLASH images, were acquired at NIH. Details for these acquisitions are described in the Supplementary information.

### Perfusion, Extraction, Cryoprotection, & Cryosectioning

Animals were perfused with 4% paraformaldehyde (PFA) through the heart, following a 50 mL saline pre-flush to remove the blood. After perfusion, the animals were decapitated posterior to the first cervical vertebrae. The heads were placed into a recirculation chamber and washed for 72 hours with 20% EDTA, followed by 5 hours in deionized water, and 1 hour in 1x PBS. The skulls were then cryoprotected in 4% PFA with 10% sucrose for 24 hours, 20% sucrose for 24 hours, and 30% sucrose for 24 hours.

The skulls were embedded in a freezing agent inside a custom negative cast mold of the block profile for each skull. The apparatus was submerged in an optimal cutting temperature compound to expedite the freezing process.

Cryosectioning of the frozen brain blocks was performed using the Microm HM550 and CryoStar NX-50 in a temperature range of -16 to -20 C and between 20% and 60% humidity. The brains were cryosectioned at 10 or 20 um using the tape transfer method^29^. Each section was placed onto a 1”x 3” slide coated in Solution B adhesive. Slides were then exposed to UV light for 8 seconds, curing the tissue onto the slide. See **Supplementary Materials: Methods** for details.

### Histological Staining

Staining was performed in alternating Nissl-Myelin or Nissl-Myelin-H&E patterns. High-throughput Nissl tissue staining was performed in an automated stainer using 1.88 g thionin in 750 mL deionized water (DiH2O), 9 mL of glacial acetic acid, and 1.08 g sodium hydroxide pellets for slide incubation. Slides then underwent three DiH2O washes and dehydration in increasing concentrations of ethanol and finally xylene before automated coverslipping.

The myelin staining technique was performed using a modified silver stain developed by Gallyas^30^. Slides were incubated in a mix of pyridine and acetic anhydride for 35 minutes, followed by a series of washes with DiH2O, EtOH, and increasing concentrations of acetic acid. The slides were immersed in a silver nitrate solution for 35 minutes, followed by the wash with 0.5% acetic acid. The slides were then developed in a mix of Developer A, Developer B and Developer C solutions. Next, the slides were washed three times with 0.5% acetic acid. Slides dried for 24 hours and were dehydrated with graded ethanol solutions before automated coverslipping.

### Microscopy image series acquisition & QC

All slides were scanned by a Nanozoomer 2.0 HT with a 20x objective (0.46 μm/pixel in plane) and saved in an uncompressed RAW format. Image cropping, conversion and compression to per section .jp2 files were performed. An online QC portal displaying high-resolution section images was used to flag damaged sections to avoid unnecessary post-processing and identify the need to repeat specific processing stages.

### Multimodal Registration Algorithms

For registration and atlas mapping, the GDM (Generative Diffeomorphic Mapping) registration algorithm is employed ^10,31^. Briefly, tissue processing procedures such as extraction and fixation cause brain tissue deformation in addition to natural biological variability, and unguided reconstruction of serial sections leads to accumulated long-range distortions ^32^. Diffeomorphic (smooth and invertible) mapping emerged to overcome these challenges. Our approach to atlas mapping and registration is a generative probabilistic model, where a synthetic stack of 2D microscopy images is formed as a sequence of transforms of a 3D image, plus a noise model describing variability. Transforms include diffeomorphic spatial warping, rigid slice positioning and contrast changes. Under this model, mapping to common coordinates is maximum a posteriori estimation, enabling the reconstruction of 3D and 2D datasets. Diffeomorphisms are estimated by gradient descent in the LDDMM framework ^33^, and linear transforms (3D affine and 2D rigid on each slice) are estimated jointly by Riemannian gradient descent as described in ^34^. We quantify tissue distortion with the derivative of spatial maps (morphometry), enabling us to account for scale changes when quantifying cell density. This tool supports multimodal registration between *in vivo* and *ex vivo* MRI, atlas annotations including independently derived segmentation, and multiple stained sections. This framework enables us to jointly analyze multiple MRI contrasts and histology data, providing ground truth data for the evaluation of MR models. In other work, we have demonstrated that MRI-constrained reconstruction shows improved accuracy over the baseline method and reduced deformable metric cost ^32^. The local scale change between postmortem MRI volumes and reassembled 3D histological volumes from the tape-transfer method is small (∼2% median absolute scale change), less than the pre-mortem to postmortem change (∼4%, measured using the Jacobian determinant of metric tensor relating the corresponding spaces)^10^.

### Creation of Atlas Volumes

To create an appropriate coordinate system defined by landmarks external to the brain, a nonlinear population average was computed from *in vivo* T2 weighted MRI, including the skull, bias corrected using the N4 algorithm ^35^. This was done using a group of six female specimens. Using a Frechet mean approach similar to the MNI152 nonlinear average ^36^ or the Allen CCF ^37^, we iteratively estimated a new average image (step 1), followed by diffeomorphic mapping from this image to each member of the population (step 2). The average image is computed by deforming each *in vivo* MRI back to common coordinates using the map computed in step 2 and performing a Jacobian-determinant weighted average at each pixel. Due to different fields of view in different samples (an issue not pertinent to the Allen CCF or the MNI space template), per-pixel weights were set to 0 for pixels outside the field of view for a given sample. Furthermore, weights were multiplied by the posterior probability that a given pixel did not correspond to an artifact or missing tissue, as described in^10^. This weighting corresponds to a maximum likelihood reconstruction. The mappings are updated using our deformable image registration tool described in the previous section. An average shape is created by minimizing the sum of square distance, in the space of diffeomorphic shape changes, from the atlas volume to each dataset^38^. A Frechet mean procedure was again used to align *ex vivo* MRI and dMRI parameter maps into the same coordinate system and to each other.

After construction of our population average image, a standard coordinate system was identified. Each of the six subjects’ MRI images was rigidly registered to a corresponding CT image. Using the CT image, the bregma point was located by creating an isosurface of the skull in the Paraview software and manually identifying a point closest to the intersection the coronal suture and sagittal suture. A symmetry plane based on the skull was identified by aligning points in the isosurface to their reflection, minimizing a measure matching loss function ^39^ over a center point and normal vector that define the symmetry plane. Once identified, the bregma point was projected onto this plane to account for any asymmetry in the sutures. Our initial estimates were found to be a root mean square distance of 0.189 mm from the symmetry plane. Points within a 2.5 mm radius sphere of this bregma position were extracted, and a normal vector to the skull was estimated by applying PCA and choosing the direction corresponding to the smallest eigenvalue of the covariance. This estimate was projected into the symmetry plane. The root mean square angle between normal vectors and before and after projection into the symmetry plane was found to be 1.87 degrees, suggesting this method is accurate but that averaging across samples is appropriate to reduce variability. The center point, the normal vector to the symmetry plane, and the normal vector to the skull together define a coordinate system for each CT scan.

A rigid transform of these coordinate systems into our *in vivo* atlas was computed by minimizing the sum of square error between voxel locations mapped by our deformable transform, and voxel locations mapped by an optimal rigid transform in the neighborhood of the bregma point, achieved with a Procrustes algorithm. Rigidly transformed coordinate systems were averaged across all six mice. The average of six bregma points was used for the origin, and the Fréchet mean ^40^ was computed for the orientations, modelling each coordinate system as a rotation matrix. The result leads to our conventions for the origin (bregma point on the skull), x axis (normal to the skull, pointing superior), y axis (left-right axis, pointing left), and z axis (anterior-posterior axis, pointing posterior). This procedure is illustrated in **Fig. 1.b**.

To produce a multimodality atlas of the brain in head, including high resolution histology (0.46 um in plane resolution) that has not been interpolated out of the sectioning plane, three representative specimens were selected for sectioning in the coronal, sagittal and transverse planes. For these individuals, deformable mappings were created from our *in vivo* average image to their Nissl series, and rigidly from the Nissl series to an interleaved myelin series using the above-described methods. The results of automatic registration were improved by manual annotations. For the Nissl series, a set of four landmark points were identified on each Nissl slice, its neighbor and its corresponding MRI section. Rigid transforms for each slice were updated by minimizing a weighted sum of square loss between Nissl and MRI, and Nissl to neighbor (encouraging smoothness). For the myelin series, a rigid transformation to the closest Nissl slice was updated using a manual procedure when the quality of the alignment was poor. We chose an additional set of three brain-only histology stacks, densely sectioned with Nissl (no interleaving myelin), in three orientations to complement the above. They were aligned using the same procedure. A lower resolution (20 um isotropic) summary dataset was also deformed backward into the same shape as our population average.

To produce initial annotations and compare to existing datasets, we also mapped our population average image to the Allen CCF^37^ and deformed their labels into our new coordinate system. We also collect cells from the Allen Brain Cell atlas ^19^ and produce xyz coordinates for each cell in our new coordinate system.

## Data Availability

Full resolution input microscopy series (Nissl and myelin) are available for viewing on the web at the Brain Architecture web portal (https://brainarchitecture.org/bap-mouse-atlas/). Reference space and analytics outputs mapped onto these microscopy series (reference space curvilinear coordinate grid and coordinate retrieval, segmentation region boundaries, Nissl cell detections, myelin segmentations) are also available as overlays on the same web viewer at the Brain Architecture web portal.

The Reference Atlas Framework is composed of a number of co-registered data volumes in the common reference space determined from skull features on the CT images co-registered with the *in vivo* MRI. These data volumes, uniformly reconstructed at 20μm isotropic resolution are all provided for download from http://data.brainarchitecture.org/. The downloadable objects are listed in Table 2 of the paper under downloadable objects.

## Code Availability

Code used to generate several main text figures, and to register individual per brain datasets with the reference space volume are made available at the following Github links:

1. https://github.com/twardlab/emlddmm.git
2. https://data.brainarchitectureproject.org/pages/mouse

## Acknowledgements

We gratefully acknowledge the generous support of the BICCN Consortium through NIH grant MH114821, the Crick-Clay Professorship and internal support at CSHL. We also recognize the significant contributions of the late Harvey Karten, whose help, advice, and insights regarding histology and neuroanatomy were invaluable to this work. Additionally, we thank Dmitry Novikov, Els Fieremans, Terezija Miskic, Xu Li and Tatiana Mitra for their assistance with various aspects of this project.

## Author contributions

C.M. initiated manuscript drafting with PPM’s guidance, created initial versions of figures and text, and was primarily responsible for correcting the annotations on the Allen reference atlas.

S.S. collected histological data for the manuscript together with P.F., M.R., B.L. J.O’R. P.F. and S.S. developed the skull decalcification protocol. S.S., P.F., M.R., B.L. and J.O’R. carried out QC, proofreading, and segmentation of the histological data.

S.B., S.Balani, K.A, contributed to the online viewer, data analysis and the portal.

R.C-L. processed MRI datasets and wrote corresponding parts of the manuscript together with J.Z.

J.Z. conducted all radiological scans for core atlas datasets, supervised R.C-L and wrote the MR components of the manuscripts.

D.T. developed algorithms for multimodal registration and average coordinate system estimation, carried out the associated tasks, and wrote the associated parts of the manuscript.

P.P.M. conceptualized and initiated the project, oversaw all project components, including experimental, computational and informatics parts, and wrote the manuscript together with the other authors.

## Materials & Correspondence

Materials and correspondence requests should be directed to. Materials made available include downloadable datasets, web-viewable outputs posted onto the Brain Architecture website, and analytic and other code repositories. Please see the **Data Availability** and **Code Availability** sections for details.

## Mouse brain atlases in the literature

We provide a list of existing mouse brain atlases in **Extended Data Table 2**. The techniques used to create these atlases vary, as do the levels of detail and resolution provided. The atlases are based on different methods such as MRI, histology, and immunohistochemistry, and are listed chronologically, with updated versions grouped together. Historically, the most widespread set of histological atlases are the stereotaxic atlases by Paxinos and Franklin, published in five editions. Many subsequent histological atlases are based on their data, making this atlas possibly the most commonly used histological print atlas of the mouse brain.

The early 2000s saw the advent of MRI mouse atlases with varying resolutions. The number of histological sections and stains varies across atlases, but none offer serial sections with the density presented in the current manuscript, nor do they simultaneously provide all three planes of section. For example, the Paxinos and Franklin atlas (2001) includes 100 coronal plates and 32 sagittal plates, while the Dong et al. (2008) atlas provides 132 coronal annotated sections in detail and 21 sagittal sections. The Paxinos and Franklin 5th edition (2019) atlas includes Nissl, AChE, and IHC for parvalbumin, consisting of 100 coronal plates, 32 sagittal plates, and 30 horizontal plates.

Some newer atlases, such as the Allen Mouse Brain Common Coordinate Framework (CCFv3) by Wang et al. (2020) and the *in vivo* MRI atlas by Meyer et al. (2017), use a combination of imaging techniques and provide more detailed and comprehensive data than earlier atlases. For example, the Allen Mouse Brain CCFv3 includes histology, immunohistochemistry, and in situ hybridization data in a 3D averaged template, while the Meyer et al. dataset includes *in vivo* MRI data with a resolution of 100 um.

In terms of limitations, some atlases may have limited applicability for certain research questions due to their focus on a specific brain regions or types of analysis. For instance, some atlases are based on a limited number of brains, while others are constrained by the resolution of the imaging techniques used. Overall, choosing the appropriate mouse brain atlas depends on the specific research question and the required level of detail and resolution. It may be necessary to consult multiple atlases or combine data from different atlases to achieve the desired level of detail and specificity.

## Supplemental section S4: MRI microgeometry parameter maps

The MD, FA, and MK maps reported in the main text illustrate typical tissue contrasts from diffusion MRI data of the mouse brain. Additional diffusion MRI parameter maps (**Extended Data** Figure 2) provide a more detailed breakdown of diffusivity and kurtosis measurements along axial and radial directions, offering specific insights into axonal and myelin injury in mouse models of neurological disorders^1^. Axial and radial directions are defined along or perpendicular to axonal bundles in white matter tracts, although these definitions become less precise in gray matter regions.

In the *in vivo* mouse brain, AD values of large white matter tracts (e.g., the corpus callosum) are higher than those in gray matter regions (e.g., the cortex), but this contrast diminishes in the *ex vivo* mouse brain. RD values of large white matter tracts are lower than those in gray matter in both *in vivo* and *ex vivo* conditions. AK and RK values in white matter tracts are higher than those in gray matter regions, with the most distinct contrasts observed in the *ex vivo* mouse brain. Overall changes in these parameters are shown using voxel-wise scatter plots in **Extended Data** Figure 3.

Beyond the empirical diffusion tensor and kurtosis models, which lack specificity to specific tissue microstructure, we incorporated parameters from a biophysical model of tissue microstructure^2^. In this and similar models^3,4^, axons are represented by long, impermeable cylinders and cell bodies by spheres. Some properties of the spatial organization of these cellular compartments can be inferred from diffusion MRI signals based on their distinct restrictive effects on water molecule diffusion. These parameters include axonal fraction (*f*), fraction of isotropic diffusion compartment (*fiso*), fiber dispersion (*p2*), intra-axonal axial diffusivity (*Da*), extracellular axial diffusivity (*D ^||^*), and extracellular radial diffusivity (*De^⊥^*). **Extended Data** Figure 2B shows representative maps of these parameters.

In the map of *f*, the difference between gray and white matter regions is more pronounced in the *ex vivo* mouse brain compared to the *in vivo* mouse brain. *fiso* is higher in the *ex vivo* corpus callosum than *in vivo.* Maps of *p2* show an increase in the *ex vivo* mouse brain in gray matter regions compared to the *in vivo* state. Changes in *Da*, *D ^||^*, and *De^⊥^* follow the same patterns as observed in the maps of AD and RD. Overall changes in these parameters are shown using voxel-wise scatter plots in **Extended Data** Figure 4.

## Supplementary Methods

### MRI & CT (Radiology)

In addition to the in vivo and ex vivo data acquired at NYU, extra CT and MRI scans were acquired at NIH:

*Ex vivo* CT (NIH): CT images of mouse heads *ex vivo* were acquired using a Quantum GX MicroCT scanner (Perkin Elmer, Hopkinton, MA, USA) in the Mouse Imaging Facility at NIH (MIF). Image acquisition time was 14 minutes per sample for nominal isotropic resolution of 32 µm.

*Ex vivo* MRI (NIH): A vertical bore 14.1 Tesla Microimaging scanner (Bruker Biospin, Billerica, MA) in the NIH MIF was used. The scan protocol included a low resolution 2D FLASH pilot, TE/TR=1.5/120 ms, Resolution = 0.156x0.156 mm^2^, Scan time = 23 s, and a high-resolution 3D FLASH TE/TR=8/30 ms, 2 averages, Resolution = 0.05x0.05x0.05 mm^3^, Scan time = 1 h 40 m.

For *in vivo* and *ex vivo* imaging, an 11.7 Tesla animal MRI system (30 cm horizontal bore magnet, Magnex Scientific, Oxford, England; MRI Electronics, Bruker Biospin, Billerica, MA) with a 12-cm integrated gradient shim system (Resonance Research Inc, Billerica, MA) was used. Animals were initially anesthetized with 3% isoflurane in 40% O2 mixed high-flow compressed air delivered at 1.5 L/min and maintained at 1% isoflurane concentration for the duration of imaging. Imaging was performed using a four-channel phased array cryogenically cooled receive-only coil with a volume transmit coil (CryoProbe system, Bruker). The scan protocol included a low resolution 2D FLASH (TE/TR=2.3/40 ms, Resolution = 0.156x0.156 mm^2^, Scan time = 1 m, 40 s) and a high resolution 3D FLASH (TE/TR=4.65/30 ms, 2 averages, Resolution = 0.05x0.05x0.05 mm^3^, Scan time = 43 m). The *ex vivo* scan protocol included the same low-resolution 2D FLASH, two repetitions of the high-resolution 3D FLASH, and an additional low-resolution 3D FLASH (TE/TR=4.65/30 ms, 2 averages, Resolution = 0.08x0.08x0.08 mm^3^, Scan time = 18 m).

### Histology & Microscopy

#### Mouse colony & facilities

All animals at CSHL were acquired from Jackson Laboratories (stock 000664) under IRB protocol #498813-18 according to protocols approved by the Animal Care and Use Committee at Cold Spring Harbor Laboratory. The animals were 56 ± 3-day-old, C5BL/6J mice, with weights ranging from 18.8 to 26.4 g. Additionally, twelve C57BL/6 mice (6 males and 6 females) were imaged at P56 and sacrificed at P56. All experimental procedures were approved by the Animal Use and Care Committee at the New York University Grossman School of Medicine.

#### Perfusion, fixation and brain extraction

The animals were sacrificed on postnatal day (P)56, and after any *in vivo* imaging experiments, by deep anesthetization with isoflurane and Avertine (2.5%). Perfusion pressure was maintained at 130 mm Hg with a Masterflex L/S Economy Pump System (Cole-Parmer, Vernon Hills, IL). The animals were perfused with 4% paraformaldehyde (PFA) (JT Baker, #JTS898-7) through the heart, after a 50 mL saline pre-flush to remove blood from the circuit. After perfusion, the animals were decapitated posterior to the first cervical vertebrae. Before imaging, the brains were stored in PBS, as well as after imaging. Heads were submerged in fomblin immediately before *ex vivo* imaging. The current imaging protocol was approximately 10-17 hours for a T1-weighted image.

#### Decalcification

After imaging, the skulls were placed in a recirculation chamber for all wash and cryoprotection steps. The skulls were washed with 20% EDTA (Sigma Aldrich, ED4S) for 72 hours, followed by dH2O for 5h, and 1x PBS for 1 hour.

#### Cryoprotection and freezing

The skulls were post-fixed for 24 hours in a solution of 4% PFA with 10% sucrose (JT Baker, #4072-05), followed by 24 hours in a solution of 4% PFA with 20% sucrose, and another 24 hours in a solution of 4% PFA with 30% sucrose. Embedding in freezing agent (Neg-50, Thermo Scientific 6505 Richard-Allan Scientific) was done using a custom-developed apparatus and a negative cast mold of the block profile for each brain. The apparatus was submerged in an optimal cutting temperature compound to expedite the freezing process.

#### Cryosectioning

Cryosectioning of the frozen skull blocks was performed using the Microm HM550 and CryoStar NX-50 in a temperature- and humidity-controlled chamber set between -16 to -20 C and 20% to 60% humidity, with the cryostat specimen head temperature set at -16 C. The cryosectioning was conducted in a temperature- and humidity-controlled room set at 18 C and 60% humidity. The skulls were coronally cryosectioned at 10 um with alternating sections using the tape transfer method ^7^. Each section was placed onto a 1”x 3” StarFrost slide first coated with Solution B adhesive. Once the appropriate amount of sections were placed onto a slide, the slides were then exposed to UV light for 8 seconds inside the cryo-chamber to cure the tissue onto the slide. The cured slides were then placed into an appropriately sized slide box and then stored at 4 C for 24 hours before starting the staining procedures.

#### Histological Staining

Separate histological staining processes were performed on different series of brain sections. High-throughput Nissl tissue staining was conducted in an automated staining machine (Sakura Tissue-Tek Prisma, DRS-Prisma-J0S). The slides underwent an automated Nissl staining protocol starting with a thionin solution: 1.88 g thionin chloride (TCI, T0214) in 750 mL deionized water (DiH2O), 9 mL glacial acetic acid (WAKO, 012–00245), and 1.08 g sodium hydroxide pellets (Sigma-Aldrich, 221465). The slides were washed three times with DiH2O, dehydrated in increasing concentrations of ethanol (50%, 70%, 95%–100%), and finally treated with xylene. The slides were then automatically cover-slipped (Sakura Tissue-tek Glas2-g2-S0) and left to dry for 24 hours.

The myelin staining technique used a modified silver stain originally developed by Gallyas^8^. The protocol was applied to the slide-mounted sections. The slides were incubated in a mix of pyridine and acetic anhydride for 35 minutes, followed by washes with DiH2O, 50% EtOH, 30% EtOH, and increasing concentrations of acetic acid. The slides were immersed in a silver nitrate solution for 35 min, followed by a wash with 0.5% acetic acid. The slides were developed using a mix of solutions: Developer A (Sodium carbonate), Developer B (Ammonium nitrate, Silver nitrate, Tungstosicilic Acid), Developer C (Ammonium nitrate, Silver nitrate, Tungstosicilic Acid 37%, Formaldehyde). The slides were washed in three consecutive washes with 0.5% acetic acid, placed on a drying rack for 24 hours, dehydrated with graded ethanol solutions, followed by automatic cover-slipping with DPX (Sigma-Aldrich). After the physical development of the myelin stain, the tissue was manually inspected for staining and morphological quality.

#### Neurohistological image series acquisition

All the prepared slides were scanned by series using a Nanozoomer 2.0 HT (Hamamatsu, Japan) with a 20x objective (0.46 μm/pixel in plane) at 8-bit depth by 3 channels and saved in an uncompressed RAW format. Imaging data were collected from the Nanozoomer and automatically transferred to a data acquisition system, which serves as the central repository for image pre-processing, including image cropping, conversion, and compression, using custom-built code in C and Matlab.

#### Quality Control

The quality control (QC) service was applied at all stages of experimentation and image data flow to correct and improve the pipeline process organically. The experimental pipeline process information was recorded in an internal Laboratory Information Management System (LIMS), which supported the workflow by recording the detailed status of each experimental stage for each brain. Similarly, a separate online QC portal managed all image pre-processing stages. Through the LIMS and QC portal, damaged sections were flagged to avoid unnecessary post-processing, and the need to repeat specific processing stages was identified.

### Downloadable Output Dataset Descriptions

Downloadable reference datasets associated with the current mouse whole head atlas resource are listed and described below:

1. *In Vivo* Averaged Reference Space Volume: This template volume is the backbone of the reference atlas resource, averaging all *in vivo* MRI from the mice used in the current manuscript. The coordinate space was defined using CT scans by fitting Bregma, Lambda, and Midline sutures, finding the intersection point between Bregma and Lambda sutures to define the origin, and defining a tangent plane orthogonal to the fit origin to define orientation. This volume is available at resolutions up to 20μm voxels, isotropic.
2. Reconstructed Nissl Volumes: Six separate Nissl volumes from six mice were reconstructed using 20μm isotropic voxels. Three datasets cover the whole head, and three includes the whole brain plus the lower brainstem and upper spinal cord. Nissl microscopy series were registered and reconstructed in reference space, with all volumes mapped to the *in vivo* reference coordinate system with Bregma as the origin.
3. Reconstructed Myelin Volumes: Three separate Myelin volumes from three mice were reconstructed using 20μm isotropic voxels. All datasets cover the whole head. Myelin microscopy series were registered and reconstructed in reference space, with all volumes mapped to the *in vivo* reference coordinate system with Bregma as the origin.
4. *In Vivo* MRI Volumes: Twelve *in vivo* volumetric MRI (T2w) datasets are upsampled and reconstructed at the resolution of the Nissl and Myelin Microscopy volumes (20μm isotropic voxels).
5. *Ex Vivo* MRI Volumes: Twelve *ex vivo* volumetric MRI (T2w) and DWI (diffusion weighted images) datasets are upsampled and reconstructed at the resolution of the Nissl and Myelin Microscopy volumes (20μm isotropic voxels).
6. Brain Region Segmentations: Brain region compartments originate from version 3 of the Allen Institute Mouse Reference Atlas but were corrected and updated as described in the manuscript. These segmentations are mapped to the reference coordinate space with Bregma as the origin and are available for download wholistically as a segmented volume (resolved at 20μm isotropic voxels), and provide a starting point for refining annotations.

### Segmented volume ID Reassignment and Integer Precision Remapping

Brain region segmented volumes used in the BAP mouse head RAF originate from version 3 of the AIBS (Dong, et al, 2020) mouse atlas. Despite having a set of approximately 700 unique region IDs, integer ID values span a range up to 2^31 in these segmented volumes, necessitating a bit precision of uint32 (32-bit unsigned integers). Additionally, homologous regions in the left and right hemispheres are not separated in the original AIBS volume (Dong, et al., 2020). To generate a unique set of region IDs for the right hemisphere and avoid conflicts with existing IDs, values equal to or greater than 2^31 were used, requiring a bit precision in the segmented volume file of uint64 (64-bit unsigned integers). This is undesirable for several reasons, including unnecessarily large file sizes and a lack of interpretability by standard segmented volume viewing packages, such as ITK-Snap (cite). Therefore, we regenerated all region IDs for the left hemisphere to range from 1 to 700 and added 2^13 to each homologous region’s ID in the right hemisphere. This allowed us to reduce file size by approximately 2 orders of magnitude and ensure interpretability by standard volume viewing packages by saving the volumes with a uint16 bit precision (16-bit unsigned integer format).

### Segmented Volume Bubble Correction

Brain region segmented volumes used in the BAP mouse head RAF originate from version 3 of the AIBS (Dong, et al, 2020) mouse atlas. Due to registration and interpolation artifacts, these volumes contain bubbles – isolated singular or small sets of voxels with region IDs that do not match their neighbors. We corrected these bubbles to eliminate artifactual isolated region ID assignment to voxels. First, we defined continuity of region IDs as voxels with the same ID assignment having touching faces. We set a threshold of five or fewer continuous voxels with the same ID assignment to constitute a bubble, or isolated set. We then identified all bubbles for each region ID and reassigned each voxel in each bubble to the same ID as the majority of its neighboring voxels not within the bubble, following a principle similar to nearest neighbor interpolation. This process was repeated until all voxels within all bubbles are reassigned. Any lingering bubbles missed by the heuristic algorithm were hand-corrected using Blender3D. This heuristic algorithm corrected approximately 99.5% of all voxels within bubbles, reducing the number of bubbles from almost 28,000 to just over 150 – a reduction of over 2 orders of magnitude. The MATLAB package for generating these volumes and the annotation volumes themselves are made available through https://data.brainarchitecture.org/pages/mouse.

---

**Extended Data Figure 1.**
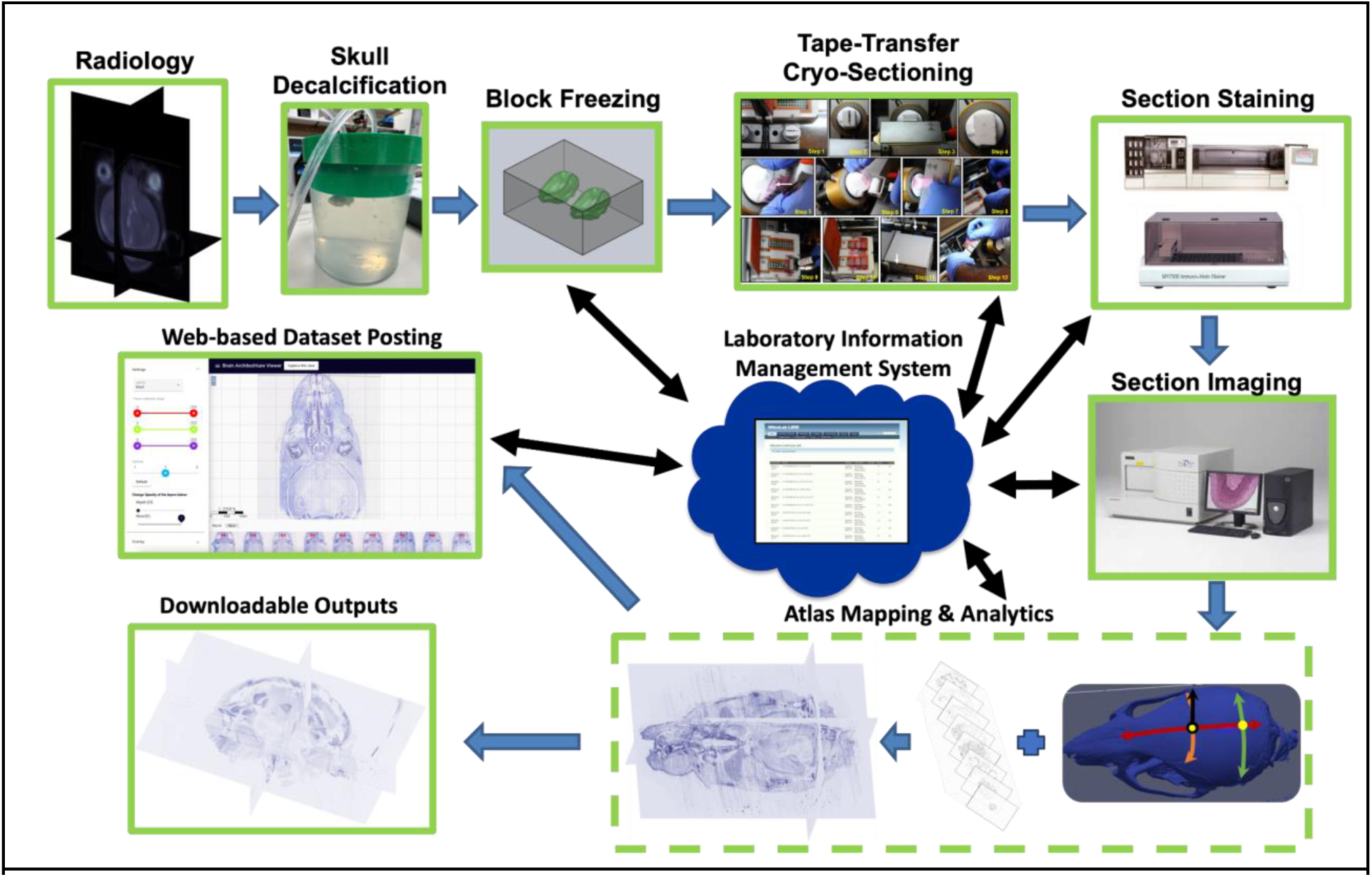
The mouse head/brain processing pipeline is summarized below. (1) Radiological images are acquired, both MRI & CT, in vivo and ex vivo. (2) Mouse heads are received at CSHL and undergo skull decalcification. (3) Mouse heads are frozen into blocks. (4) Blocks are tape-transfer cryosectioned. (5) Sections are stained on slide using an automated stainer for both Nissl and Myelin histology. (6) Sections are imaged with a Hamamatsu Nanozoomer. (7) Following section image QC, section images are aligned and mapped into the in vivo stereotactic reference coordinate space. (8) Following additional analytics and atlas mapping, per subject outputs are released in both web viewable and downloadable formats. Black arrows indicate steps with key integration with the LIMS (an in-house built Laboratory Information Management System).

**Extended data Figure 2.**
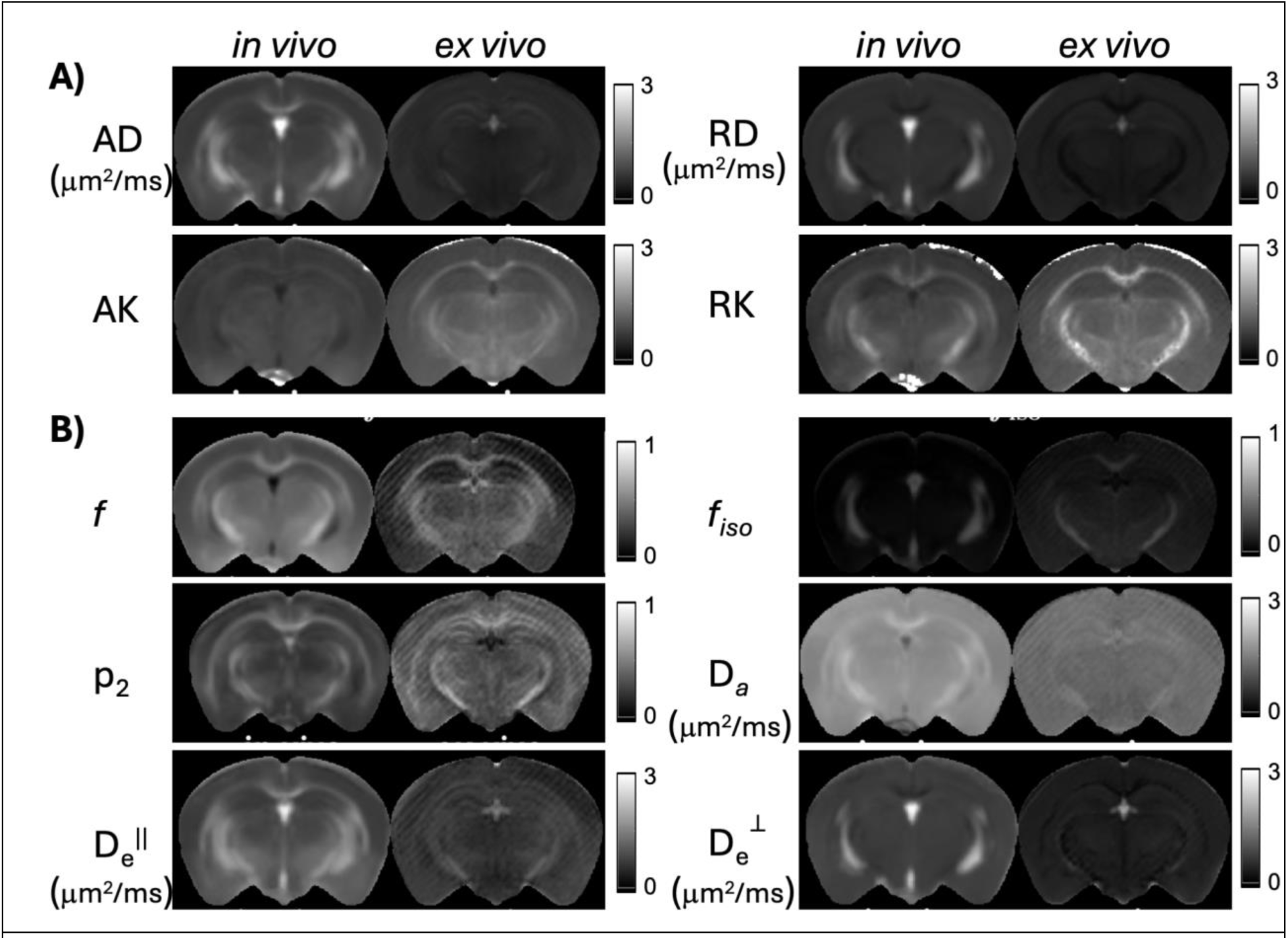
Additional *in vivo* and *ex vivo* population-averaged diffusion MRI parameter maps of the female mouse brain. **A:** *In vivo* and *ex vivo* axial diffusivity (AD), radial diffusivity (RD), axial kurtosis (AK), and radial kurtosis (RK) derived from the empirical diffusion tensor and kurtosis model. **B:** *In* vivo and *ex vivo* tissue microstructure parameters derived from a biophysical model, in which axons are approximated by long impermeable cylinders and cell bodies by spheres. Abbreviations: *f* : axonal fraction; *fiso*: fraction of isotropic diffusion compartment; p2: fiber dispersion; Da: intra-axonal axial diffusivity; De^||^: extracellular axial diffusivity; De^⊥^ : extracellular radial diffusivity.

**Extended Data Figure 3.**
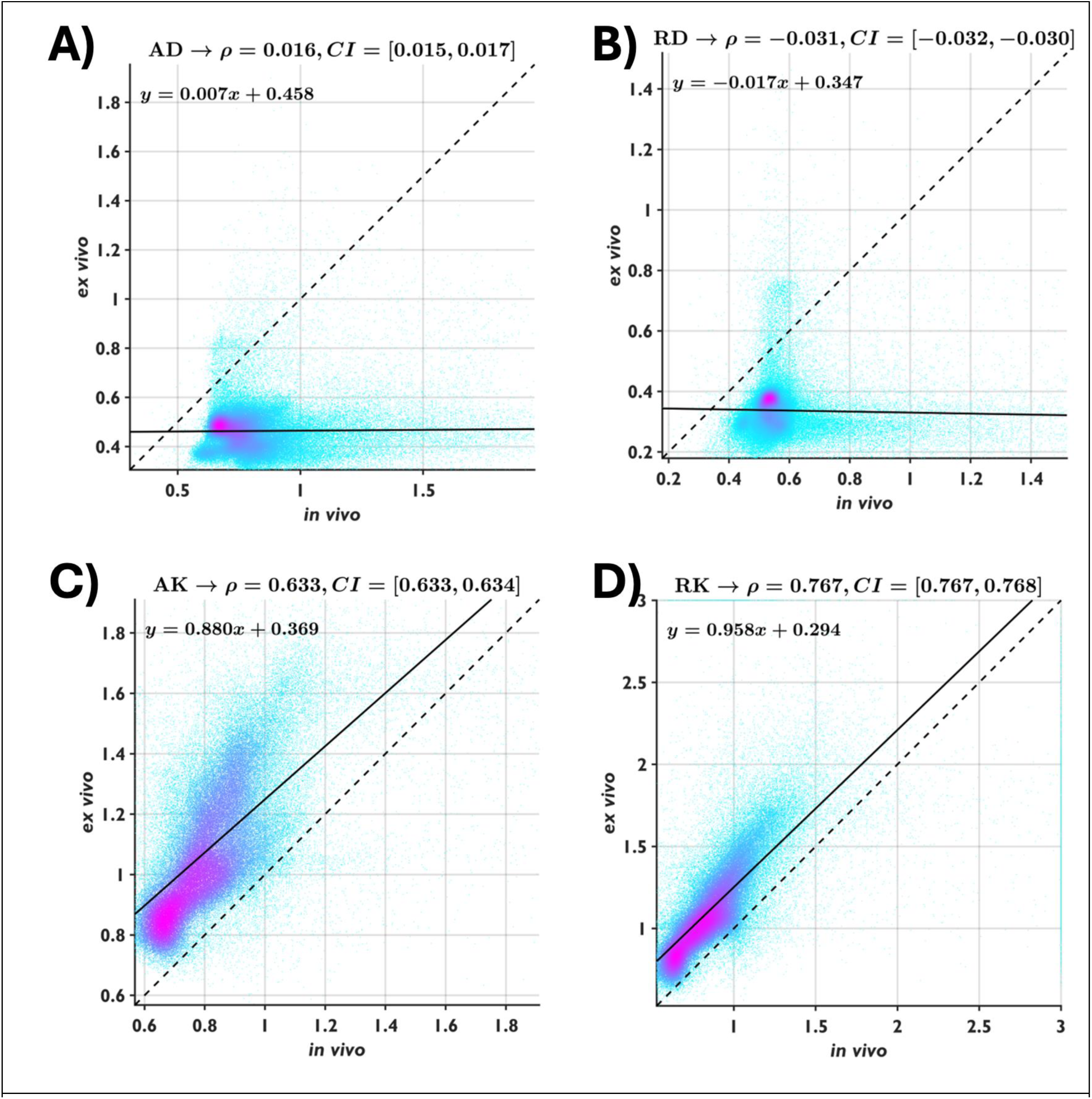
Voxel-wise comparisons between *in vivo* and *ex vivo* values of axial diffusivity (AD), radial diffusivity (RD), axial kurtosis (AK), and radial kurtosis (RK), derived from the empirical diffusion tensor and kurtosis model and shown in scatter plots with Pearson correlation coefficient (𝜌) and 95% confidence interval (CI). The solid lines represent the results of linear regression, and the dashed diagonal lines indicate the reference line for no change. The color indicates the density of voxels. Both *ex vivo* AD and RD are reduced compared to *in vivo* AD and RD, whereas *ex vivo* AK and RK show slight increases compared to *in vivo* AK and RK.

**Extended Data Figure 4.**
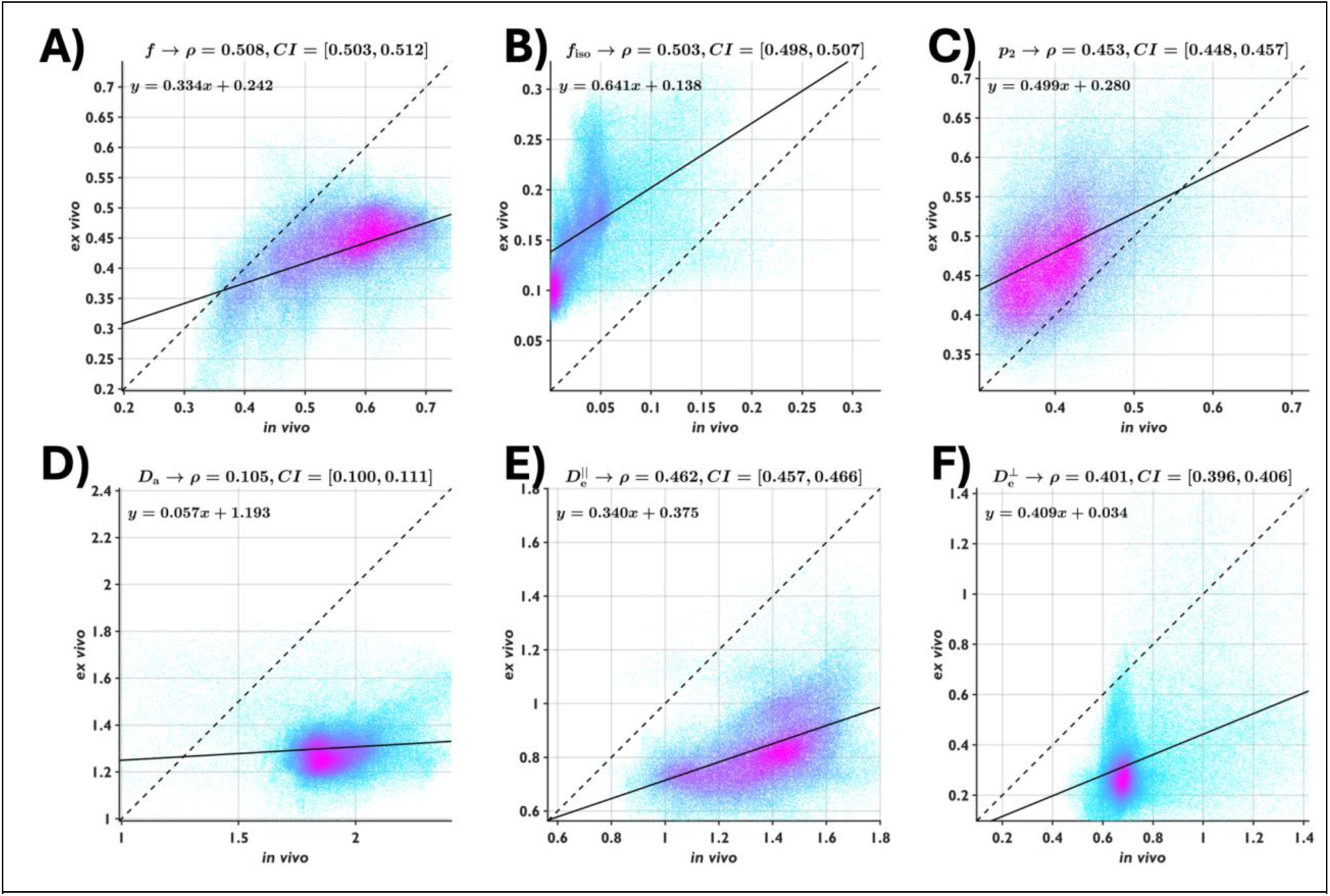
Voxel-wise comparisons between *in vivo* and *ex vivo* values of axonal fraction (*f*), fraction of isotropic diffusion compartment (*f_iso_*), fiber dispersion (p_2_), intra-axonal axial diffusivity (D_a_), extracellular axial diffusivity (D_e_^||^), and extracellular radial diffusivity (D_e_^⊥^), computed from the biophysical model and shown in scatter plots with Pearson correlation coefficient (𝜌) and 95% confidence interval (CI). The solid lines represent the results of linear regression, and the dashed diagonal lines indicate the reference line for no change. *Ex vivo f*, D_a_, D_e_^||^, and D_e_^⊥^ are lower than the *in vivo* results, and ex vivo *f_iso_* and p_2_ are higher than the *in vivo* results.

**Extended Data Figure 5.**
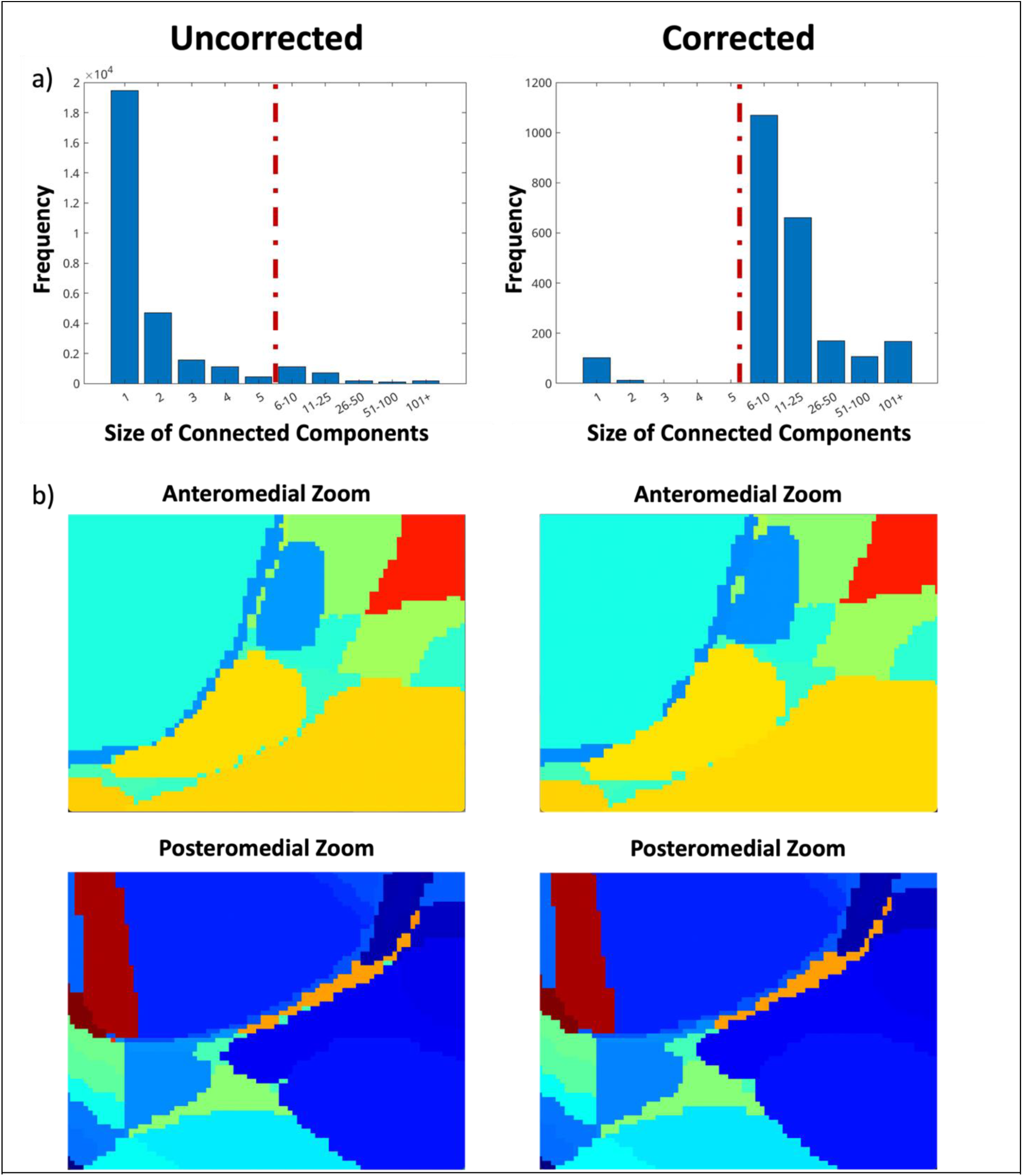
Demonstration of output of bubble correction in action. (a) Histograms of the size of connected components for both the uncorrected and corrected volumes. Note the dashed red line after a voxel size of 5, defining all components whose size was less than 5 voxels as a bubble. (b) Anteromedial and posteriomedial zoom in panels showing the segmentations at a per-voxel resolution both before and after bubble correction. Note the reduction in isolated voxels in the right, corrected anteromedial and posteromedial panels relative to the left uncorrected anteromedial and posteromedial panels.

**Extended Data Figure 6.**
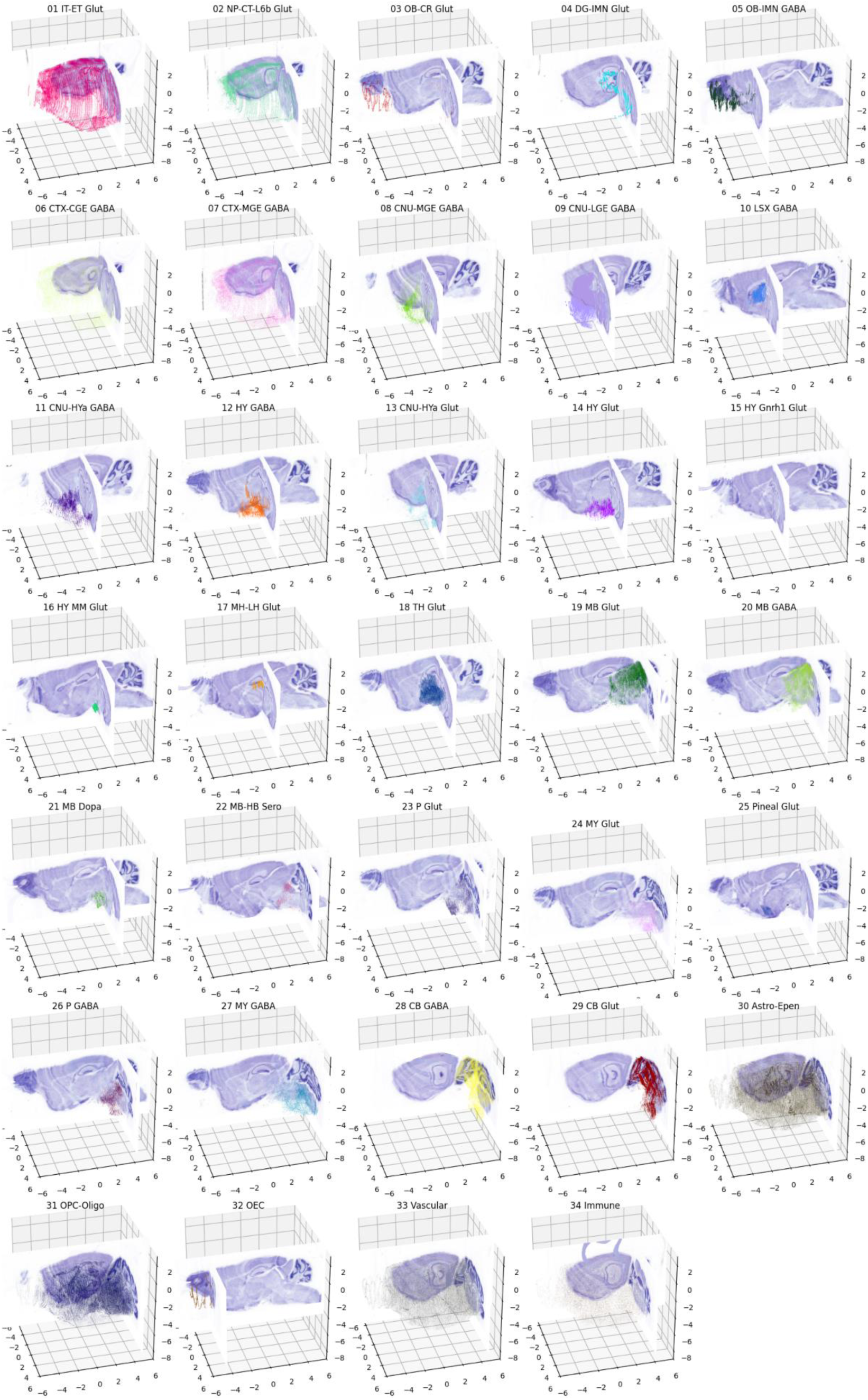
Distribution of 35 cell types from the ABC atlas in our reference atlas coordinate system. Here we show the distribution of 35 different cell types loaded using the Allen Brain Cell (ABC) Atlas python API as described at the url https://alleninstitute.github.io/abc_atlas_access/notebooks/merfish_ccf_registration_tutorial.html, with coordinates mapped from the Allen CCF into our reference atlas framework. Sagittal and coronal sections from one of our Nissl volumes are shown such that 90% of cells are rendered in front of the section. Coordinates are in mm, with the bregma point at (0,0,0). Cells are rendered as a 3D scatter plot, with colors using the conventions from the API.

**Extended Data Table 1.** Subject metadata and histology series (P56, C5BL/6J mice)

| Subject ID | Sex | Section Plane | Skull | Radiology | Histology |
| --- | --- | --- | --- | --- | --- |
| MD961 | F | Coronal | + | In vivo, ex vivo MRIs, ex vivo CT | Nissl, myelin, 10µm alternate sections |
| MD962 | F | Transverse | + | In vivo, ex vivo MRIs, ex vivo CT | Nissl, myelin, 10µm alternate sections |
| MD963 | F | Sagittal | + | In vivo, ex vivo MRIs, ex vivo CT | Nissl, myelin, 10µm alternate sections |
| MD585 | M | Sagittal | - | N/A | Nissl, 20µm serial sections |
| MD634 | M | Coronal | - | N/A | Nissl, 20µm serial sections |
| MD636 | M | Transverse | - | N/A | Nissl, 20µm serial sections |

**Extended Data Table 2.**
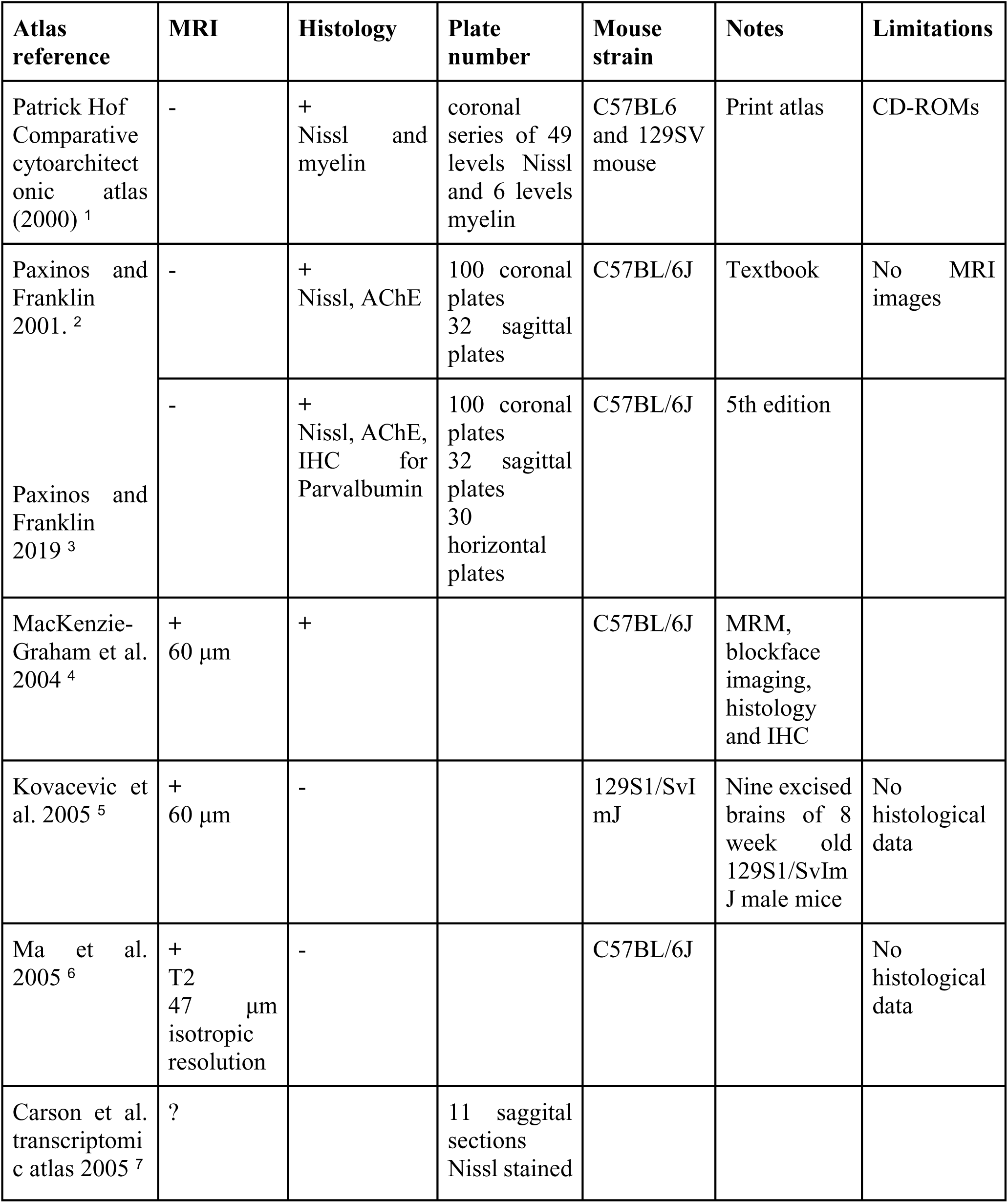
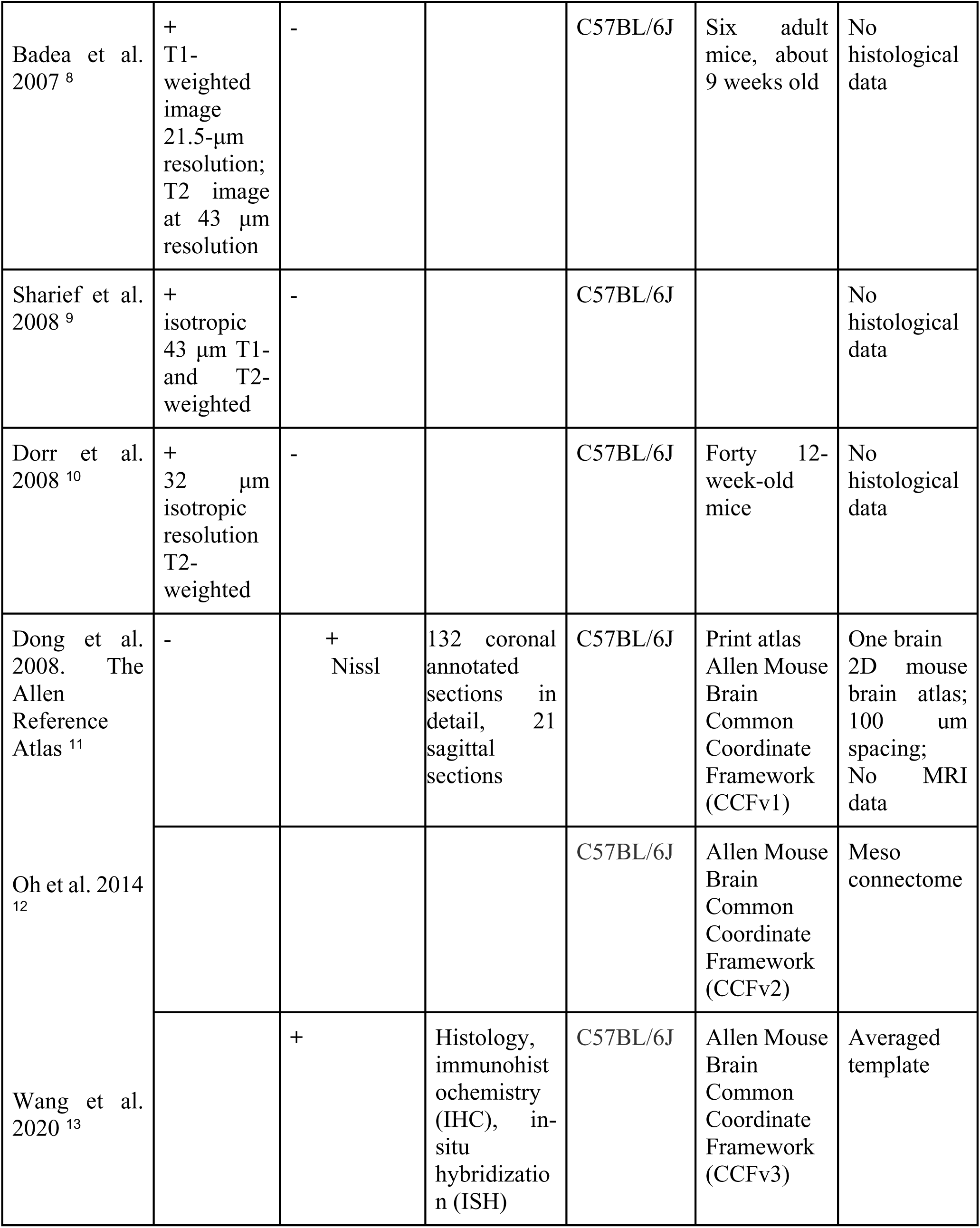
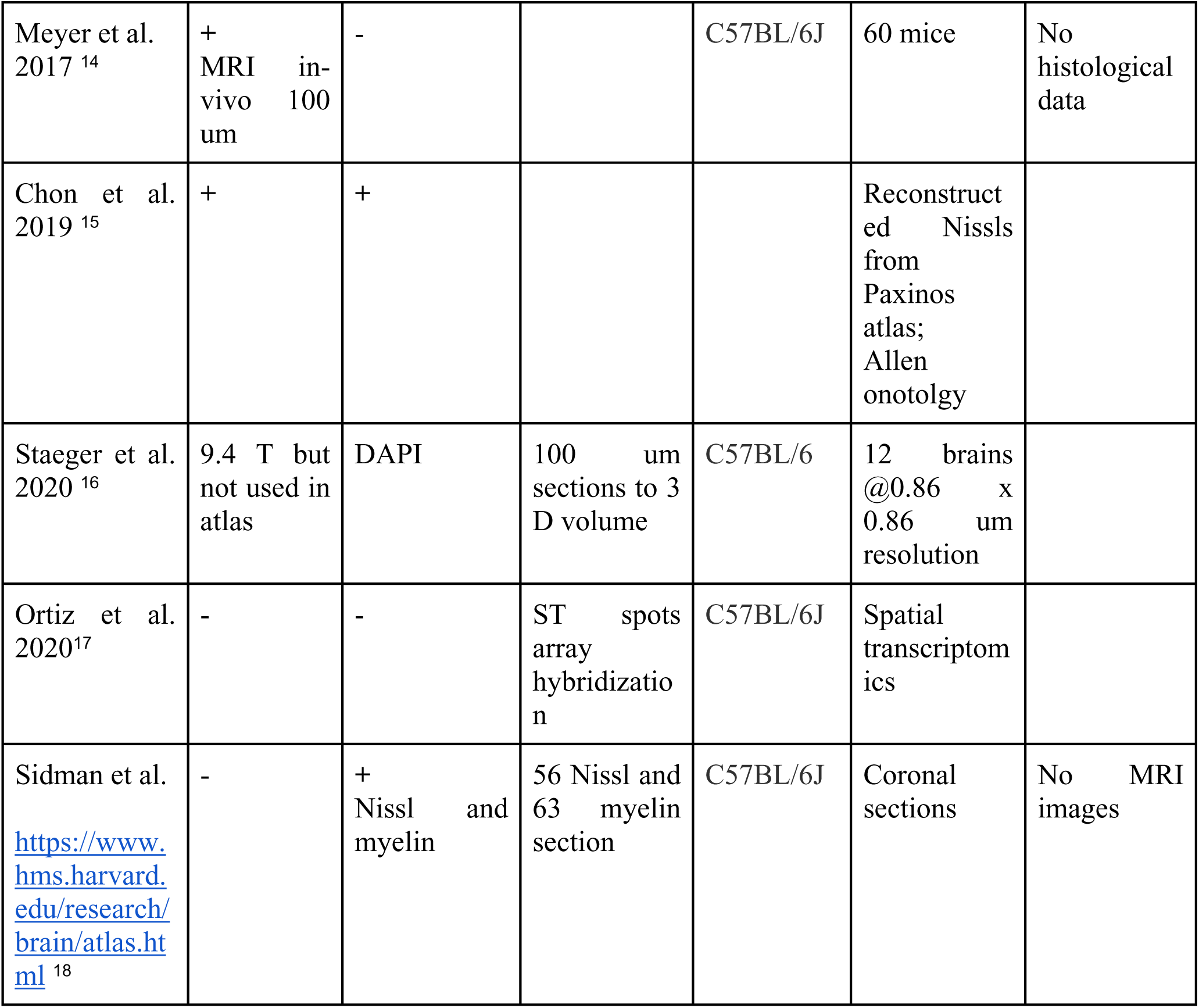
Current mouse brain atlases with histological data.

**Extended Data Table 3.** Existing MRI atlases of the mouse brain.

| Author (s) | Mouse strain(s) | Imaging modalities | Brain regions covered | # of structures | Sampling Resolution ($\mu\text{m}^3$) | In or Ex Vivo | MRI contrast | Ages |
| --- | --- | --- | --- | --- | --- | --- | --- | --- |
| MacKenzie-Graham et al., 2003<sup>19</sup> | C57BL/6 | MRI and histology | Whole brain | 70 | 60 | ex vivo | Diffusion weighted | P100 |
| MacKenzie-Graham et al., 2004<sup>4</sup> | C57BL/6 | MRI and histology | Whole brain | 70 | 60 | ex vivo | Diffusion weighted | P100 |
| Kovacevic et al., 2005<sup>5</sup> | 129SI/SvIm | MRI | Whole brain | 42 | 60 | ex vivo | T2-weighted | P63 |
| Ma et al., 2005<sup>6</sup> | C57BL/6 | MRI | Whole brain | 20 | 47 | ex vivo | T2-weighted, diffusion tensor MRI | P84-P98 |
| Lee et al., 2012<sup>20</sup> | C57BL/6 | MRI | Whole brain | 13 | 40 | ex vivo | T2-weighted | P0 |
| Ali et al., 2005<sup>21</sup> | C57BL/6 | MR | Whole brain | 21 | 90 | ex vivo | T2-weighted | P63 |
| Chen et al., 2006<sup>22</sup> | C57BL/6, 129SI/SvIm, CD1 | MR | Whole brain | 42 | 60 | ex vivo | T2-weighted | P56 |
| Bock et al., 2006<sup>23</sup> | cdf/cdf, cdf/+ | MR and histology | Cerebrum & cerebellum | 5 | 156 | in vivo | T1-weighted | P77 |
| Johnson et al., 2007<sup>24</sup> | C57BL/6 | MR | Whole brain | 33 | 43 | ex vivo | T1, T2-weighted | P63-84 |
| Dorr et al., 2007<sup>25</sup> | CBA | MR and CT | Whole brain & vasculature | 26 | 32 | ex vivo | T2-weighted | P84 |
| Badea et al., 2007 a & b <sup>8</sup> | C57BL/6 | MR | Whole brain | 33 | 43 | ex vivo | T1, T2-weighted | P63 |
| Sharief et al., 2008<sup>9</sup> | C57BL/6 | MR | Whole brain | 33 | 43 | ex vivo | T1, T2-weighted | P63 |
|  | And BXD |  |  |  |  |  |  |  |
| Ma et al., 2008<sup>26</sup> | C57BL/6 | MR | whole brain | 20 | 100 | in vivo | T2-weighted | P84-P98 |
| Dorr et al., 2008<sup>10</sup> | C57BL/6 | MR | Whole brain | 62 | 32 | ex vivo | T2-weighted | P84 |
| Aggarwal et al., 2009<sup>27</sup> | C57BL/6 | MR and CT | Whole brain | 62 | 32 | ex vivo, in vivo for P140-160 | T2-weighted, diffusion tensor MRI | P7, P14, P21, P28, P63, P140-160 |
| Richards et al., 2011<sup>28</sup> | C57BL/6 | MR | Hippocampus | 40 | 30 | ex vivo | T2*-weighted | P84 |
| Chuang et al., 2012<sup>29</sup> | C57BL/6 | MR | Developing brain (cortex, hippocampus, cerebellum, and white matter tracts) | 62 | 80 to 125 | ex vivo | T2-weighted, diffusion tensor MRI | E12-P80 |
| Ullmann et al., 2013<sup>30</sup> | C57BL/6 | MR | Neocortex | 74 | 15 | ex vivo | T2*-weighted | P84 |
| Steadman et al., 2014<sup>31</sup> | NL3 KI | MR | Cerebellum | 39 | 56 | ex vivo | T2-weighted | NA |
| Szulc et al., 2015<sup>32</sup> | FVB/N | MR | Developing brain (whole brain) |  | 100 | in vivo | T1-weighted | P1-P11 |
| Kronman et al., 2023<sup>33</sup> | C57BL/6 | MR and 3D Light Microscopy | Developing brain (whole brain) | 270 | 100 | ex vivo | T2-weighted, diffusion tensor MRI | E11-P56 |

